# O-GlcNAc transferase regulates glioblastoma acetate metabolism via regulation of CDK5-dependent ACSS2 phosphorylation

**DOI:** 10.1101/2021.05.10.443157

**Authors:** Lorela Ciraku, Zachary A. Bacigalupa, Jing Ju, Rebecca A. Moeller, Rusia H. Lee, Michael D. Smith, Christina M. Ferrer, Sophie Trefely, Mary T. Doan, Wiktoria A. Gocal, Luca D’Agostino, Wenyin Shi, Joshua G. Jackson, Christos D. Katsetos, Nathaniel W. Snyder, Mauricio J. Reginato

## Abstract

Glioblastomas (GBMs) preferentially generate acetyl-CoA from acetate as a fuel source to promote tumor growth. O-GlcNAcylation has been shown to be elevated by increasing O-GlcNAc transferase (OGT) in many cancers and reduced O-GlcNAcylation can block cancer growth. Here, we identify a novel mechanism whereby OGT regulates acetate-dependent acetyl-CoA production by regulating phosphorylation of acetyl-CoA synthetase 2 (ACSS2) by cyclin-dependent kinase 5 (CDK5). OGT is required and sufficient for GBM cell growth and regulates acetate conversion to acetyl-CoA. Elevating O-GlcNAcylation in GBM cells increases phosphorylation of ACSS2 on Ser-267 in a CDK5-dependent manner. Importantly, we show that ACSS2 Ser-267 phosphorylation regulates its stability by reducing polyubiquitination and degradation. ACSS2 Ser-267 is critical for OGT-mediated GBM growth as overexpression of ACSS2 Ser-267 phospho-mimetic rescues growth *in vitro* and *in* vivo. Importantly, we show that pharmacologically targeting OGT and CDK5 reduces GBM growth *ex vivo*. Thus, the OGT/CDK5/ACSS2 pathway may be a way to target altered metabolic dependencies in brain tumors.

## Introduction

Glioblastoma (GBM) is the most common type of primary malignant brain tumor (Dunn et al., 2012). Glioblastomas, along with brain metastasis, oxidize acetate via the metabolic enzyme acetyl-CoA synthetase short chain family member 2 (ACSS2), which catalyzes the ATP-dependent conversion of acetate and coenzyme A (CoA) into acetyl-CoA (Schug et al., 2016). ACSS2 is required for the majority of acetate conversion to acetyl-CoA (Comerford et al., 2014) which can be utilized for *de novo* biosynthesis of fatty acids and to provide carbon for the TCA cycle which is essential for tumor growth and survival (Lewis et al., 2015; Mashimo et al., 2014) (Comerford et al., 2014). Reducing ACSS2 levels can block cancer cell growth both *in vitro* and *in vivo* (Comerford et al., 2014; Mashimo et al., 2014; Yoshii et al., 2009). While ACSS2 is emerging as a novel therapeutic target for brain cancer, little is known regarding its regulation.

The hexosamine biosynthetic pathway utilizes major metabolites to generate (Marshall et al., 1991) UDP-N-acetylglucosamine (UDP-GlcNAc) which serves as a substrate for both N- and O-Glycosylation that regulate cellular behaviors in response to nutrient availability (Zachara and Hart, 2004). UDP-GlcNAc is also a substrate for O-linked β-N-acetylglucosamine (O-GlcNAc) transferase (OGT) which catalyzes the addition of O-GlcNAc moieties onto serine and threonine residues of nuclear and cytoplasmic proteins. Removal of O-GlcNAcylation is catalyzed by the glycoside hydrolase O-GlcNAcase (OGA) (Gao et al., 2001). This modification alters protein functions, can regulate protein-protein interactions and phosphorylation states (Zachara and Hart, 2006) (Bond and Hanover, 2015). OGT and O-GlcNAcylation are elevated in most cancers (Ferrer et al., 2016), and targeting this modification inhibits tumor cell growth *in vitro* and *in vivo* (Caldwell et al., 2010) (Lynch et al., 2012). Recently, metabolomic profiling of breast cancer cells depleted of OGT revealed significant changes in lipid metabolites (Sodi et al., 2018). Thus, we hypothesized that OGT may regulate lipid-dependent cancers including glioblastomas.

Cyclin-dependent kinase 5 (CDK5) is an atypical member of the cyclin-dependent kinase family predominantly expressed in the brain and activated by non-cyclin activators such as p35, or its truncated product p25 (Pozo and Bibb, 2016). While CDK5 is not known to be mutated in cancers (Pozo and Bibb, 2016), deregulation of CDK5 stimulates hyperactivation of downstream proteins such as RB, STAT3, FAK, vimentin (Antoniou et al., 2011; Fu et al., 2004; Futatsugi et al., 2012; Lindqvist et al., 2015; Pozo and Bibb, 2016; Sharma et al., 2020) and regulates tumorigenesis through cell proliferation and metastatic invasion (Lenjisa et al., 2017; Pozo and Bibb, 2016). While CDK5 levels have been found to be elevated in glioblastoma tissue (Catania et al., 2001; Liu et al., 2008; Yushan et al., 2015), its functional roles in this cancer are not known.

Here, we present evidence that OGT and O-GlcNAcylation levels are elevated in glioblastoma tissues and cells and that OGT is required for GBM cell growth *in vitro* and *in vivo*. OGT regulates acetate metabolism via regulation of ACSS2 protein levels. Mechanistically, we show that increased OGT activity results in CDK5-mediated ACSS2 phosphorylation on Ser-267, stabilizing ACSS2 protein levels, and reducing polyubiquitination and degradation. OGT regulation of ACSS2 requires CDK5 and its effect on GBM growth requires Ser-267 phosphorylation of ACSS2. Importantly, ACSS2 Ser-267 phosphorylation is elevated in human glioblastoma and pharmacologically targeting OGT or CDK5 reduces GBM tumors *ex vivo* suggesting that this pathway is required for GBM growth and may serve as potential therapeutic targets for treating brain cancer.

## Results

### OGT and O-GlcNAc levels are elevated in glioblastoma and are required for tumor growth *in vitro* and *in vivo*

OGT and O-GlcNAcylation have been shown to be elevated in multiple cancers (Ferrer et al., 2016). However, the expression and possible role of OGT in glioblastomas has not been investigated. We first examined levels of OGT from normal glial tissue through progressing stages of glioma including glioblastoma (Grade IV). Using immunohistochemistry (IHC) analysis, we observed an increase in OGT-positive staining that correlates with disease progression (Fig. 1a). Notably, normal glial tissue exhibits minimal OGT-positive staining compared to Grade IV glioma (Fig. 1a). Moreover, analysis of 69 Grade IV glioblastoma patient tissue samples for global O-GlcNAcylation via IHC showed 82% of these samples expressed medium to high O-GlcNAc (Fig. 1b). We also found that primary GBM cells contain elevated OGT and O-GlcNAcylation compared to normal human astrocytes (Fig. 1c). Similar elevation of OGT and O-GlcNAcylation was detected in established GBM U87-MG and T98G cell lines (Fig. 1d). Together, these results show that human glioblastomas contain elevated levels of OGT and O-GlcNAcylation.

**Fig 1.**
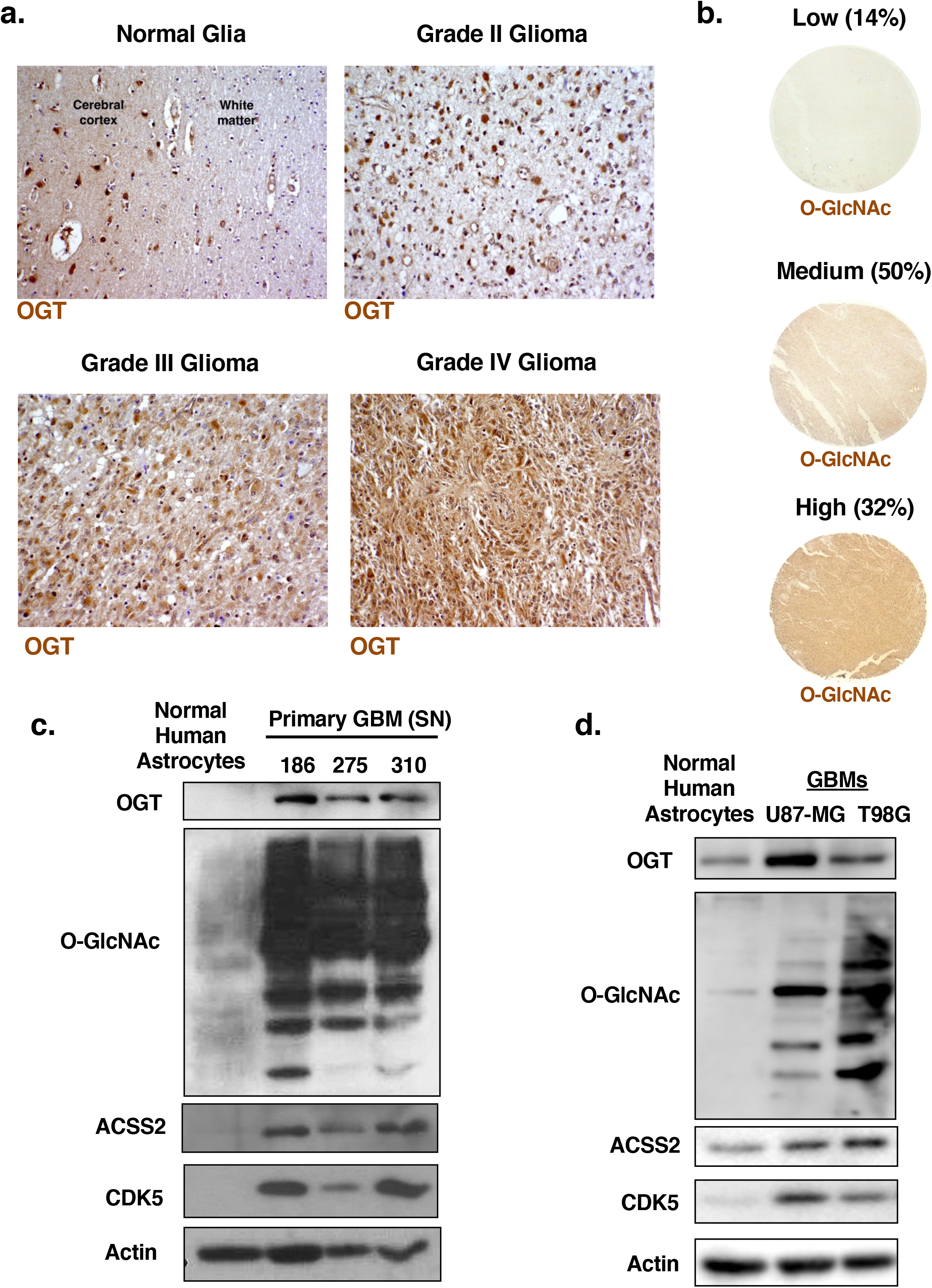
OGT and O-GlcNAcylation levels are elevated in human glioblastoma. **(a)** Immunohistochemical staining for OGT in different grades of glioma. **(b)** Immunohistochemical staining for O-GlcNAc on a tissue microarray (n=69) of Grade IV glioblastoma patient biopsies (4X scale bar 1000 um). **(c)** Cell lysate from normal human astrocytes and primary cells isolated from three different GBM patients were collected for immunoblot analysis with the indicated antibodies. **(d)** Cell lysates of normal human astrocytes, U87-MG and T98G glioblastoma cell lines were collected for immunoblot analysis with the indicated antibodies.

To examine whether OGT is required for glioblastoma growth, we targeted OGT with shRNA in glioblastoma cell lines. We observed that suppression of OGT was sufficient to impede cell growth in both U87-MG (Fig. 2a, 2b) and T98G (Fig. 2c, 2d) cells, as indicated by crystal violet staining. We also tested whether altering OGT expression could impact anchorage-independent growth. Reducing OGT expression with stable expression of RNAi (Fig. 2a, 2c), we observed a significant reduction in the anchorage-independent growth in both U87-MG and T98G (Fig. 2e, Extended Data Fig. 1a) compared to controls. Additionally, we found that reducing OGT levels in two different primary GBM cell lines SN310 and SN186 (Fig. 2f) significantly blocked neurosphere formation (Fig. 2g, 2h). To ensure OGT contribution to GBM growth required its catalytic function, we also examined effects of treating GBM cells with a pharmacological inhibitor of O-GlcNAcylation Ac-5S-GlcNAc (Ferrer et al., 2014; Gloster et al., 2011; Sodi et al., 2015). Treating U87-MG cells with Ac-5S-GlcNAc reduced total O-GlcNAcylation and also reduced cell growth (Extended Data Fig. 1b). To test whether reducing OGT expression could block GBM cell growth *in vivo*, we utilized an intracranial orthotopic xenograft model. Reduction of OGT expression in U87-MG cells expressing luciferase (Extended Data Fig. 1c) resulted in a significant decrease in tumor growth *in vivo* as measured by bioluminescence (Fig. 2i). Representative images of H&E sections show a striking reduction in tumor growth at Day 21 in mice injected with OGT depleted cells (Fig. 2i). Importantly, mice that were grafted with glioblastoma cells where OGT expression was reduced also had significantly extended survival (Fig. 2j), displaying a median survival of 42 days, compared to 20 days for the control group. Altogether, these data indicate that OGT and O-GlcNAc levels are elevated in human glioblastoma and OGT is required for glioblastoma cell growth *in vitro* and *in vivo*.

**Fig 2.**
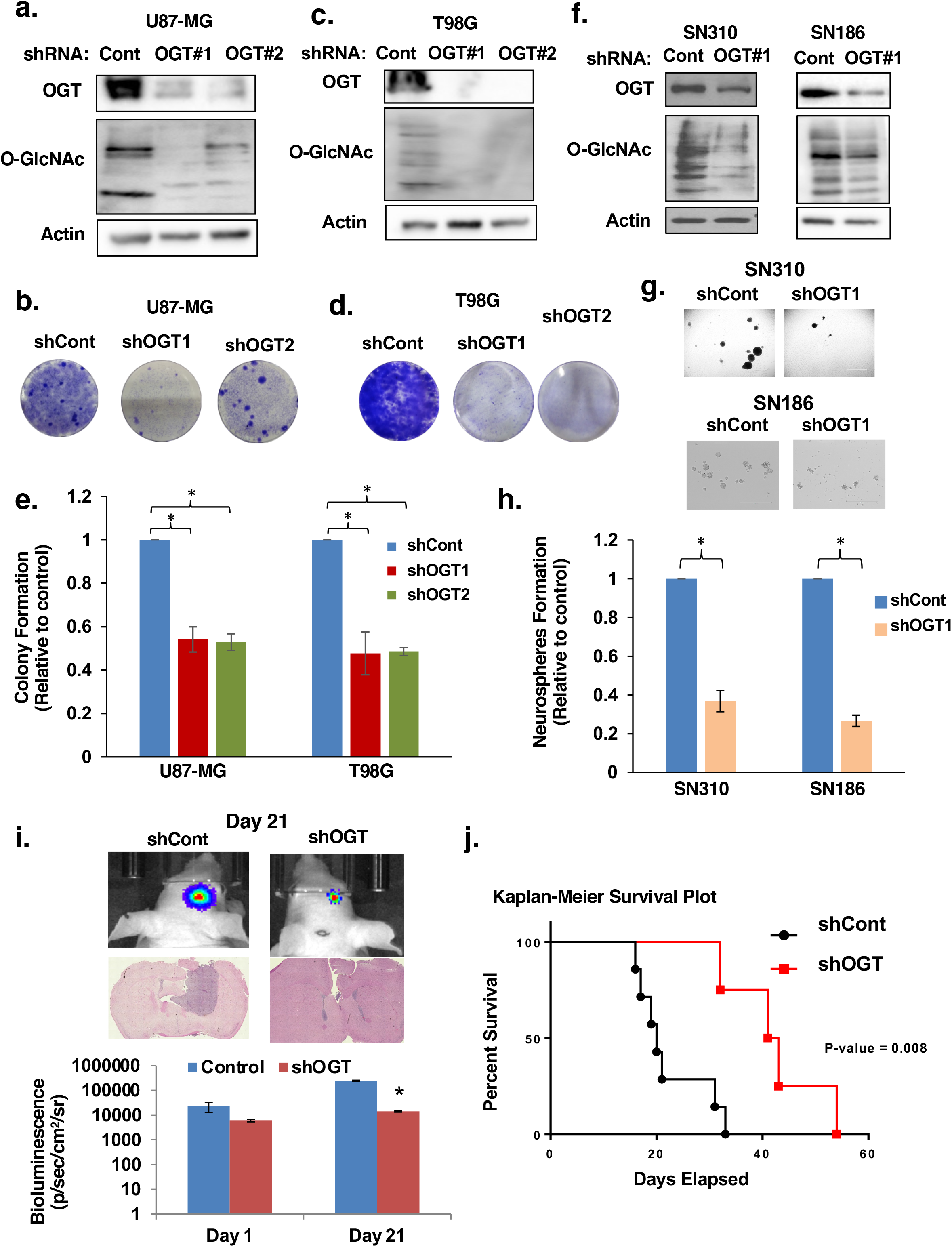
OGT is required for glioblastoma growth *in vitro* and *in vivo*. **(a)** Cell lysates from U87-MG cells expressing control or OGT shRNA were collected for immunoblot analysis with the indicated antibodies. **(b)** U87-MG cells infected with shControl or shOGT lentivirus effect on cell growth visualized with crystal violet staining. **(c)** Cell lysates from T98G cells expressing control or OGT shRNA were collected for immunoblot analysis with the indicated antibodies. **(d)** T98G cells infected with shControl or shOGT lentivirus effect on cell growth visualized with crystal violet staining. **(e)** Anchorage-independent growth assay comparing the colony formation of control or OGT shRNA expressing in indicated cells. Data are quantified and presented as average relative to control from three independent experiments. Student’s t-test reported as mean ± SEM; *p<0.05. **(f)** Cell lysates from SN310 (left) or SN186 (right) primary GBM cells expressing control or OGT shRNA where collected for immunoblot analysis with the indicated antibodies. **(g)** Cells from (f) were then plated for a neurosphere formation assay for 8 days and representative image taken. **(h)** Neurospheres were quantified for SN310 and SN186 cells and presented as average relative to control from three independent experiments. Student’s t-test reported as mean ± SEM. * = p-value < 0.05. **(i)** Representative images of bioluminescent (top) detection of tumors from mice injected with shControl and shOGT U87-MG cells 21 days post-injection. Representative images of H&E analysis (middle) on coronal sections from mice harboring shControl or shOGT tumors at Day 21. Data are quantified and presented as average from mice injected with U87-MG cells expressing shControl (n=5) or shOGT mice (n=7). (bottom). Student’s t-test reported as mean ± SEM; *p<0.005. **(j)** Kaplan-Meier survival plot comparing overall survival of mice inoculated with shControl (n=7) or shOGT (n=4) U87-MG cells. Mantel-Cox log rank test P = 0.008.

### OGT regulates lipid accumulation, acetyl-CoA, and acetate metabolism

Several groups have characterized the importance of lipid metabolism (Lewis et al., 2015), and more specifically, acetate metabolism for supporting glioblastoma growth (Comerford et al., 2014; Mashimo et al., 2014). Thus, we investigated the effect of altering OGT on lipid accumulation, and acetate dependent acetyl-CoA metabolism in glioblastoma cells. Consistent with OGT being critical for GBM cell growth, overexpression of OGT in U87-MG cells increased total O-GlcNAcylation (Fig. 3a) and anchorage-independent growth (Fig. 3b and Extended Data Fig. 1d). OGT overexpressing GBM cells also contained increased intracellular lipid droplet accumulation as measured by nile red staining (Fig. 3c), increased free fatty acids (Fig. 3d) and cellular acetyl-CoA levels (Fig. 3e). Conversely, stable knockdown of OGT in U87-MG (Extended Data Fig. 2a) cells contained reduced nile red staining (Extended Data Fig. 2b), free fatty acids (Extended Data Fig. 2c) and acetyl-CoA (Extended Data Fig. 2d) levels compared to control cells. Similar results were seen in T98G cells containing stable knockdown of OGT (Extended Data Fig. 2e-h) while treating T98G cells with OGA inhibitor increased acetyl-CoA levels (Extended Data Fig. 2i).

**Fig 3.**
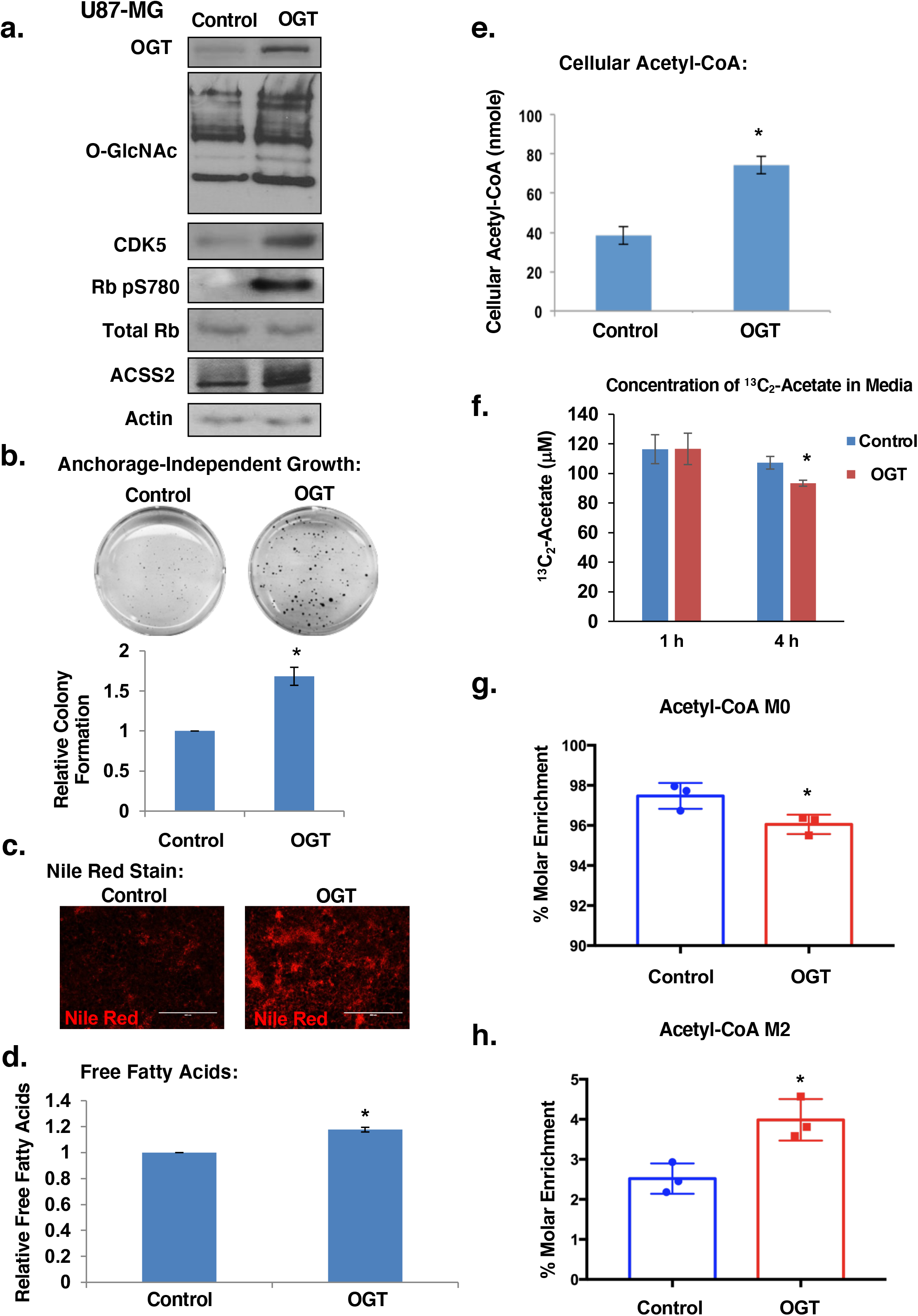
OGT promotes acetate and lipid accumulation in glioblastoma cells. **(a)** Cell lysates from U87-MG cells stably overexpressing control or OGT were collected for immunoblot analysis with the indicated antibodies. **(b)** Control or stably overexpressing OGT U87-MG cells were placed in soft agar assay. Representative images from anchorage-independent growth assay (top) and colonies were counted and quantified (bottom). Student’s t-test reported as mean ± SEM. * = p-value < 0.005. **(c)** Representative images of nile red staining of U87-MG cells under same conditions as in (a). **(d)** Measurement of relative free fatty acids in U87-MG cells stably overexpressing control or OGT. Student’s t-test reported as mean ± SEM. * = p-value < 0.05. **(e)** Measurement of acetyl-CoA extracted from U87-MG cells stably overexpressing control or OGT. Student’s t-test reported as mean ± SEM. * = p-value < 0.05. **(f)** Graphical representation of the concentration of ^13^C_2_-acetate remaining in the media following 1 and 4 hours of exposure to 100μM ^13^C_2_-acetate supplemented serum-free media in U87-MG cells stably overexpressing control or OGT. Student’s t-test reported as mean ± SEM. * = p-value < 0.05. **(g)** Graphical representation of the percent molar enrichment of unlabeled acetyl-CoA in U87-MG cells stably overexpressing control or OGT for 4 hours. Student’s t-test reported as mean ± SEM. * = p-value < 0.05. **(h)** U87-MG control and OGT overexpressing cells were labeled with 100 μM ^13^C_2_-acetate for 4 hours. Measurement of the percent molar enrichment for labeled (M2) acetyl-CoA. Student’s t-test reported as mean ± SEM. * = p-value < 0.05.

Since glioblastoma cells are highly dependent on acetate metabolism to generate acetyl-CoA and regulate growth (Mashimo et al., 2014), we hypothesized that OGT may regulate acetate utilization. To test this, we exposed U87-MG control cells and cells overexpressing OGT with stable isotope labeled sodium acetate (1,2-^13^C_2_-acetate) to track the incorporation of acetate into acetyl-CoA. Cells overexpressing OGT displayed significantly increased acetate uptake as labeled acetate was depleted from the media (Fig. 3f). Moreover, within the context of the increased pool of acetyl-CoA, we detect a 2% reduction in molar enrichment of unlabeled acetyl-CoA M+0 (Fig. 3g) and a corresponding increase in molar enrichment of acetyl-CoA M2 (Fig. 3h) in OGT overexpressing cells compared to control cells indicating an increased utilization of acetate in generating acetyl-CoA. These data implicate a role for OGT in regulating the metabolism of acetate into acetyl-CoA and contributing to GBM lipid accumulation and growth.

### O-GlcNAcylation regulates ACSS2 S267 phosphorylation in a CDK5-dependent manner

ACSS2 is a critical regulator of acetate conversion to acetyl-CoA and for growth of glioblastoma cells (Comerford et al., 2014; Schug et al., 2015). Therefore, we examined ACSS2 expression in the context of altered O-GlcNAcylation. Consistent with previous findings, we found ACSS2 levels to be elevated in both primary GBM (Fig. 1c) and established GBM cell lines (Fig. 1d) compared to normal astrocytes. U87-MG cells stably expressing OGT shRNA contained lower levels of ACSS2 protein (Extended Data Fig. 2a). RNA levels of ACSS2 were not reduced in OGT depleted U87-MG cells as measured by quantitative RT-PCR (qRT-PCR) (Extended Data Fig. 3a). Similar decreases in ACSS2 protein levels were seen in OGT depleted primary GBM cells SN310 and SN186 (Fig. 4a) and T98G (Extended Data Fig. 2e) cells. Conversely, we observe stabilization of ACSS2 protein when U87-MG cells (Fig. 3a and Extended Data Fig. 3b) or primary GBM SN310 cells (Extended Data Fig. 3c) were stably overexpressing OGT or U87-MG cells were treated with an OGA inhibitor NButGT (Extended Data Fig. 3d). ACSS2 protein does not appear to be directly modified by O-GlcNAc, as immunoprecipitated ACSS2 from normal human astrocytes and GBM cells U87-MG, T98G and SN310 contained no detectable O-GlcNAcylation by western blot analysis (data not shown). Thus, O-GlcNAcylation likely regulates ACSS2 protein stability via an indirect mechanism.

**Fig 4.**
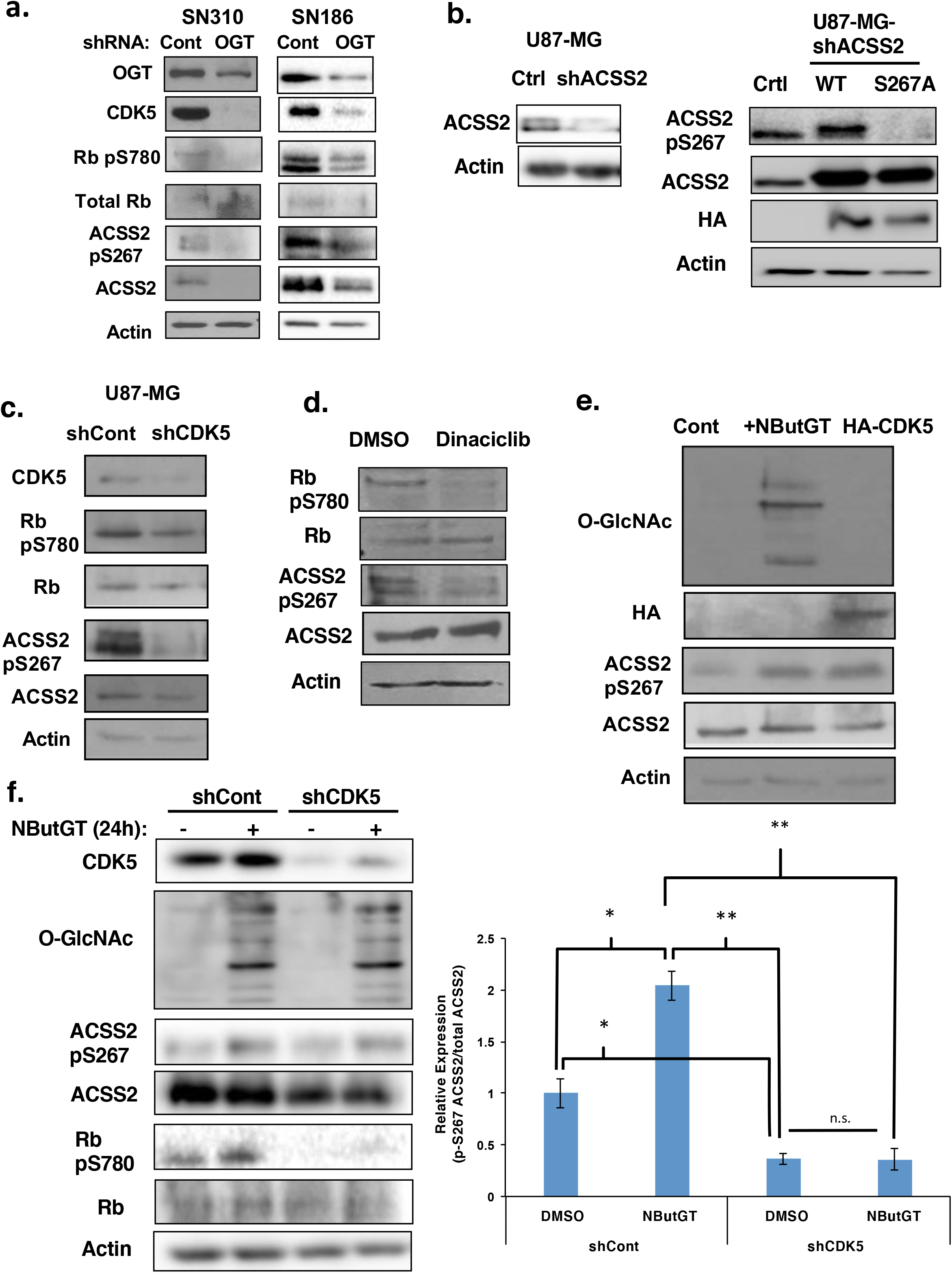
Ser267-ACSS2 phosphorylation is mediated by O-GlcNAcylation in a CDK5-dependent manner. **a)** Cell lysates from SN310 (left) or SN186 (right) primary GBM cells expressing control or OGT shRNA were collected for immunoblot analysis with the indicated antibodies. **(b)** Cell lysates from U87-MG cells stably expressing shRNA against endogenous ACSS2 and overexpressing control, wildtype ACSS2 or ACSS2 S267A mutant were collected for immunoblot analysis with the indicated antibodies. **(c)** Cell lysates from of U87-MG cells stably expressing shRNA against control or CDK5 were collected for immunoblot analysis with the indicated antibodies. **(d)** Cell lysates from U87-MG cells treated for 48 hrs with pan-CDK inhibitor dinaciclib (5 nM) were collected for immunoblot analysis with the indicated antibodies. **(e)** Cell lysate from U87-MG cells that were transfected with control plasmid with and without treatment of 100 μM OGA inhibitor (NButGT) for 24 hours or HA-CDK5 plasmid were collected for immunoblot analysis with the indicated antibodies. **(f)** Cell lysate from U87-MG cells stably expressing control or CDK5 shRNA and treated with DMSO or 100 μM NButGT for 24 hours were collected for immunoblot analysis with the indicated antibodies (left). Relative expression of p-S267 ACSS2/total ACSS2 was quantified (right). Student’s t-test reported as mean ± SEM. * = p-value < 0.05.

O-GlcNAcylation can alter the phosphorylation state of proteins (Bond and Hanover, 2015). To analyze potential changes in ACSS2 phosphorylation regulated by O-GlcNAcylation, we analyzed changes in exogenous immunoprecipitated ACSS2 phosphorylation via mass-spectrometry in GBM cells treated with control or OGA inhibitor (Extended Data Fig. 3e). Proteomic analysis identified only Ser-267 of ACSS2 as being phosphorylated in NButGT-treated conditions (data not shown). Predictive analyses for phosphorylation sites (ScanSite; (Obenauer et al., 2003)) for ACSS2 revealed Ser-267 as a putative CDK5 phosphorylation site (Extended Data Fig. 3f). Alignment analysis revealed the consensus CDK5 motif containing Ser-267 (SQ**S**PPIKR) as being evolutionarily conserved in mammals (Extended Data Fig. 3g). Consistent with our data, PhosphoSitePlus (Hornbeck et al., 2015) lists 13 studies that have identified human ACSS2 Ser-267 as being phosphorylated in different cancers including breast, lung, cervical, ovarian, and skin cancers. However, the functional role of this phosphorylation is not known. Therefore, these data suggest that elevated O-GlcNAcylation increases ACSS2 phosphorylation on Ser-267, a putative CDK5 site.

To establish ACSS2 as a putative substrate for CDK5 phosphorylation, we performed an *in vitro* kinase assay. We observed a protein concentration-dependent increase in ACSS2 phosphorylation by CDK5 (Extended Data Fig. 3h), confirming ACSS2 as a bona-fide substrate of CDK5. Importantly, ACSS2 Ser-267 to alanine (S267A) mutant contained reduced phosphorylation by CDK5 *in vitro* (Extended Data Fig. 3i). To test whether Ser-267 was also phosphorylated in GBM cells, we knocked down endogenous ACSS2 with stable expression of shRNA against ACSS2 targeting the 3’ UTR and overexpressed wildtype ACSS2 or ACSS2 S267A mutant. Mutation of ACSS2 S267 into alanine abrogated ACSS2 phosphorylation, which was detected using an antibody specifically recognizing ACSS2 pS267 (Fig. 4b). To further confirm the specificity of the phospho-ACSS2-S267 antibody, we immunoprecipitated wild-type ACSS2 and ACSS2-S267A mutant from U87-MG cells and found that the ACSS2 p267 antibody only detected wild-type ACSS2 protein (Extended Data Fig. 3j).

Phosphorylation of ACSS2 on Ser-267 in glioblastoma cells is dependent on CDK5 activity as stable expression of CDK5 shRNA reduces phosphorylation of ACSS2 on Ser-267 (Fig. 4c) as well as phosphorylation of RB on Ser-780 (Fig. 4c), a CDK5 specific phosphorylation site (Lin et al., 2007; Piedrahita et al., 2010). Similar results were seen in cells treated with the pan-CDK inhibitor dinaciclib that targets CDK5 (Fig. 4d) (Parry et al., 2010b). Conversely, overexpression of HA-CDK5 in U87-MG cells increased ACSS2 Ser-267 phosphorylation (Fig. 4e). In addition, we found that CDK5 and ACSS2 interact *in vitro* (Extended Data Fig. 3k). However, in GBM cells this interaction was only detected in conditions of elevated O-GlcNAcylation as immunoprecipitation of CDK5 showed an interaction with ACSS2 only in cells treated with OGA inhibitor (Extended Data Fig. 3l). CDK5 was also found to interact with phosphorylated ACSS2-S267, determined by immunoprecipitating with antibody specifically recognizing ACSS2 pS267, and this interaction was increased under conditions of elevated O-GlcNAcylation (Extended Data Fig. 4a). Consistent with the mass spectrometry results, we detected an increase of ACSS2 Ser-267 phosphorylation in cells treated with an OGA inhibitor (Fig. 4e and 4f). Conversely, inhibition of O-GlcNAcylation by treating cells with OGT inhibitor (Ac-5S-GlcNAc) reduced ACSS2 Ser-267 phosphorylation (Extended Data Fig. 4b). We also tested whether OGT-mediated phosphorylation of ACSS2 occurred in normal brain cells or tissue. Increased O-GlcNAcylation in normal human astrocytes treated with an OGA inhibitor did not alter ACSS2 Ser-267 phosphorylation compared to GBM cells (Extended Data Fig. 4c). Using an *ex vivo* brain slice model (Jackson et al., 2014), treating tumor-free brain tissue with an OGA inhibitor elevated O-GlcNAcylation but had no effect on ACSS2 Ser-267 phosphorylation (Extended Data Fig. 4d) suggesting that OGT-dependent regulation of ACSS2 Ser-267 phosphorylation may be cancer cell-specific. Increased O-GlcNAcylation also increased CDK5 activity as measured by phosphorylation of RB on Ser-780 (Fig. 4f). Importantly, O-GlcNAc regulation of ACSS2 Ser-267 phosphorylation was reduced in cells stably expressing CDK5 shRNA (Fig. 4f). Consistent with these results, cells overexpressing OGT contain increased CDK5 levels and activity (Fig. 3a) and GBM cells stably expressing OGT RNAi contain reduced levels of CDK5 and activity and ACSS2 ser-267 phosphorylation (Fig. 4a, Extended Data Fig. 2a, 2e). These results indicate that O-GlcNAc regulates ACSS2 phosphorylation on Ser-267 in a CDK5-dependent manner.

### ACSS2 S267 phosphorylation blocks degradation of ACSS2 and is required for GBM growth

Since we observed stabilization of ACSS2 protein when GBM cells were overexpressing OGT (Fig. 3a, Extended Data Fig. 3b, 3c) or following OGA inhibition (Extended Data Fig. 3d), we hypothesized that OGT and CDK5 regulate ACSS2 stability via phosphorylation of Ser-267. To test whether OGT regulates ACSS2 degradation, U87-MG cells stably expressing control shRNA or OGT shRNA were treated with the proteasome inhibitor lactacystin. OGT shRNA-mediated inhibition of ACSS2 protein levels could be reversed by treatment with lactacystin (Fig. 5a) suggesting ACSS2 is being regulated at the proteasomal level. To test whether ACSS2 Ser-267 phosphorylation alters polyubiquitination of ACSS2, U87-MG cells stably expressing wild-type ACSS2 and S267 phosphosite mutants were transfected with ubiquitin (WT-Ub) or ubiquitin-K48R (K48R-Ub) mutant. Following immunoprecipitation ofACSS2, we show that wild-type ACSS2 is polyubiquitinated and that ACSS2-S267A mutant contains significantly increased polyubiquitination compared to wild-type ACSS2 and ACSS2-S267D phospho-mimetic mutant in cells transfected with WT-Ub (Fig. 5b, Extended Data Fig. 4e). Since mutation of lysine 48 to arginine renders ubiquitin (Ub) unable to form poly-Ub chains via lysine 48 linkages with other Ub molecules (Chau et al., 1989), we tested effect of K48R-Ub on polyubiquitination of ACSS2-S267A. K48 polyubiquitination of ACSS2-S267A was significantly reduced in cells transfected with ubiquitin K48R mutant (K48R-Ub) compared to WT-Ub (Fig. 5b, Extended Data Fig. 4e). This data suggests that Ser-267 phosphorylation alters polyubiquitination of ACSS2. Consistent with the idea that ACSS2 phosphorylation of Ser-267 stabilizes and blocks degradation of ACSS2, we show that the protein half-life of ACSS2 S267A mutant is significantly reduced compared to wildtype ACSS2 or ACSS2 S267D phospho-mimetic mutant (Fig. 5c) in cells treated with cyclohexamide. To test whether phosphorylation of ACSS2 Ser-267 plays a role in tumorigenesis in GBM cells, we stably reduced endogenous ACSS2 expression utilizing a shRNA targeting the 3’UTR (Fig. 5d) and then stably re-expressed either wildtype ACSS2, ACSS2 S267A or ACSS2 S267D phospho-mimetic mutant (Fig. 5d). U87-MG cells overexpressing wildtype ACSS2 or phosphomimetic S267D mutant, but not ACSS2 S267A mutant, were able to rescue anchorage independent growth (Fig. 5e). Consistent with results seen *in vitro*, U87-MG cells expressing wild-type

**Fig 5.**
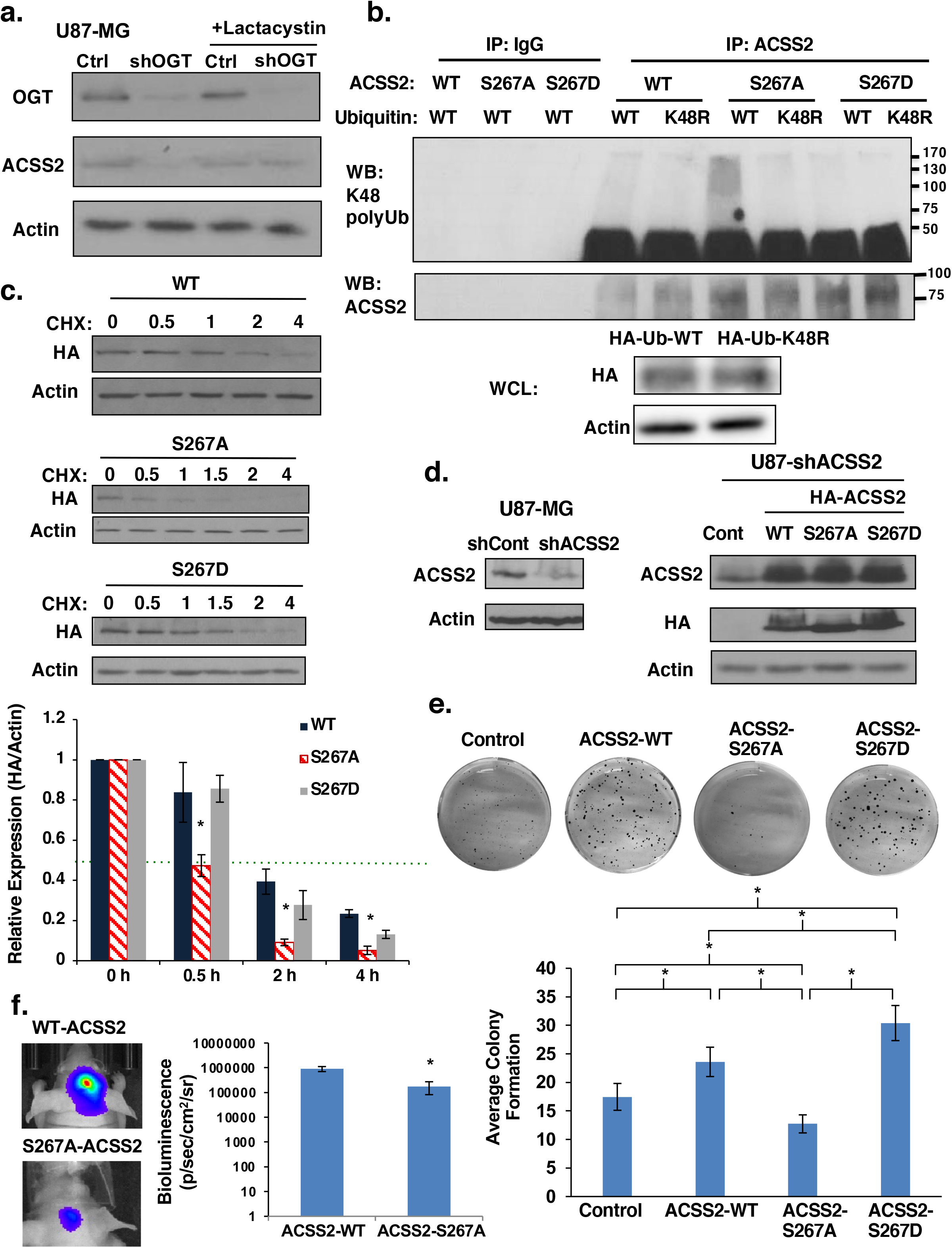
Phosphorylation of Ser267 enhances stability of ACSS2 and is required for GBM growth. **(a)** Cell lysate from U87-MG cells stably expressing control or OGT shRNA and treated with the proteasomal inhibitor 10 μM lactacystin for 6 hours were collected for immunoblot analysis with the indicated antibodies. **(b)** Immunoprecipitation was performed with the indicated antibodies from U87-MG cell lysates stably expressing wild-type (WT)-, S267A-, or S267D-HA-ACSS2 and transfected with Ub-WT or Ub-K48. **(c)** Cell lysate from U87-MG cells stably expressing wild-type (WT)-, S267A-, or S267D-HA-ACSS2 in U87-MG glioblastoma cells treated with 10 μg/μl cycloheximide for indicated time (hours) were collected for immunoblot analysis with the indicated antibodies (top). Densitometry quantification of three independent time-course experiments presented as relative expression (HA/actin) to 0 hours (bottom). Green dotted line represents 0.5 relative expression. Student’s t-test reported as mean ± SEM. * = p-value < 0.05. **(d)** Cell lysates from U87-MG cells stably expressing ACSS2 shRNA against endogenous 3’ UTR of ACSS2 (left) and stably overexpressing wildtype ACSS2, ACSS2-S267A and ACSS2-S267D mutant (right) were collected for immunoblot analysis with the indicated antibodies. **(e)** Representative image of cells in (d) seeded into an anchorage-independent growth assay and imaged at day 14 (top). Data are quantified and presented as average from at least three independent experiments (bottom). Student’s t-test reported as mean ± SEM. * = p-value < 0.05. **(f)** Representative images of tumor growth detected via bioluminescence at Day 16 following injection of U87-MG-luciferase cells WT-ACSS2 or S267A-ACSS2 (left). Quantification of tumor size (WT n=4, S267A n=4) (right). Student’s t-test reported as mean ± SEM. * = p-value < 0.05

ACSS2, but not cells containing ACSS2 S267A mutant, were able to form tumors *in vivo* (Fig. 5f, Extended Data Fig. 4f). These results indicate that phosphorylation of ACSS2 on S267 is critical for its protein stability, polyubiquitination and for GBM cell growth *in vitro* and *in vivo*.

### CDK5 regulates acetate metabolism and GBM growth via ACSS2-S267 phosphorylation

CDK5 has been implicated in the development and progression of multiple cancers (Pozo and Bibb, 2016). However, its role in glioblastoma has not been demonstrated. CDK5 levels have been shown to be elevated in GBM tissue (Liu et al., 2008). A high level of CDK5 expression predicted poor prognosis in GBM patients (Extended Data Fig. 5a). Consistent with this data, we detect an increase in CDK5 levels in primary (Fig. 1c) and established GBM cell lines (Fig. 1d) compared to normal human astrocytes. Since CDK5 phosphorylates ACSS2 S267 *in vitro* and in cells, we tested the role of CDK5 on acetate metabolism and GBM cell growth. To test whether CDK5 regulates acetate metabolism and GBM cell growth we reduced CDK5 levels via shRNA which blocked CDK5 activity as measured by RB phosphorylation on Ser-780 and reduced ACSS2 phosphorylation on Ser-267 (Fig. 6a). We observed that suppression of CDK5 was sufficient to impede the growth of U87-MG cells as indicated by crystal violet staining (Fig. 6b) and anchorage-independent growth (Extended Data Fig. 5b). Reducing CDK5 levels in T98G cells also blocked growth (Extended Data Fig. 5c) and anchorage-independent growth (Extended Data Fig. 5d) and knockdown in primary GBM cells significantly blocked neurosphere formation (Fig. 6c, Extended Data Fig. 5e). Consistent with the idea that CDK5 regulates ACSS2, we also found that depleting CDK5 levels in GBM cells reduced acetyl-CoA levels (Fig. 6d) and nile red staining (Extended Data Fig. 5f). Importantly, we show that reduction in CDK5 expression in U87-MG cells resulted in a significant decrease of 1,2-^13^C_2_-acetate incorporation into acetyl-CoA (Fig. 6e). Moreover, we show that reduction of CDK5 expression in U87-MG cells resulted in a significant decrease in tumor growth *in vivo* as measured by bioluminescence and histology (Fig. 6f). To test whether OGT and O-GlcNAc-mediated growth in GBM cells was dependent on CDK5 expression we tested the effect of reducing CDK5 in OGT overexpressing cells. The increase in anchorage-independent growth observed in U87-MG cells stably overexpressing OGT was reduced as a result of CDK5 knockdown cells (Extended Data Fig. 6a, 6b). Consistent with OGT regulating ACSS2, the observed OGT-mediated increase in ACSS2-S267 phosphorylation (Extended Data Fig. 6c) and acetyl-CoA (Extended Data Fig. 6d) was reduced in CDK5 knockdown cells. We detected similar decrease of ACSS2-S267 phosphorylation (Extended Data Fig. 6e) and acetyl-CoA (Extended Data Fig. 6f) in U87-MG cells treated with OGA inhibitor containing CDK5 knockdown. To test whether ACSS2 phosphorylation was required for CDK5 depletion-mediated effects, we overexpressed wildtype HA-ACSS2, HA-ACSS2 S267A and HA-ACSS2 S267D mutant in the context of CDK5 knockdown (Fig. 6g). Cells stably overexpressing exogenous wildtype ACSS2 and ACSS2 S267D mutant, but not ACSS2 S267A mutant, partly restored growth (Fig. 6h) and nile red staining (Extended Data Fig. 6g) in CDK5 knockdown cells. Thus, these data suggest that CDK5 can regulate acetate metabolism and GBM growth *in vitro* and *in vivo* and that OGT-mediated cell growth and acetyl-CoA regulation is partly dependent on CDK5. In addition, we show that CDK5-mediated growth in glioblastoma cells is partly dependent on phosphorylation of ACSS2 Ser267.

**Fig 6.**
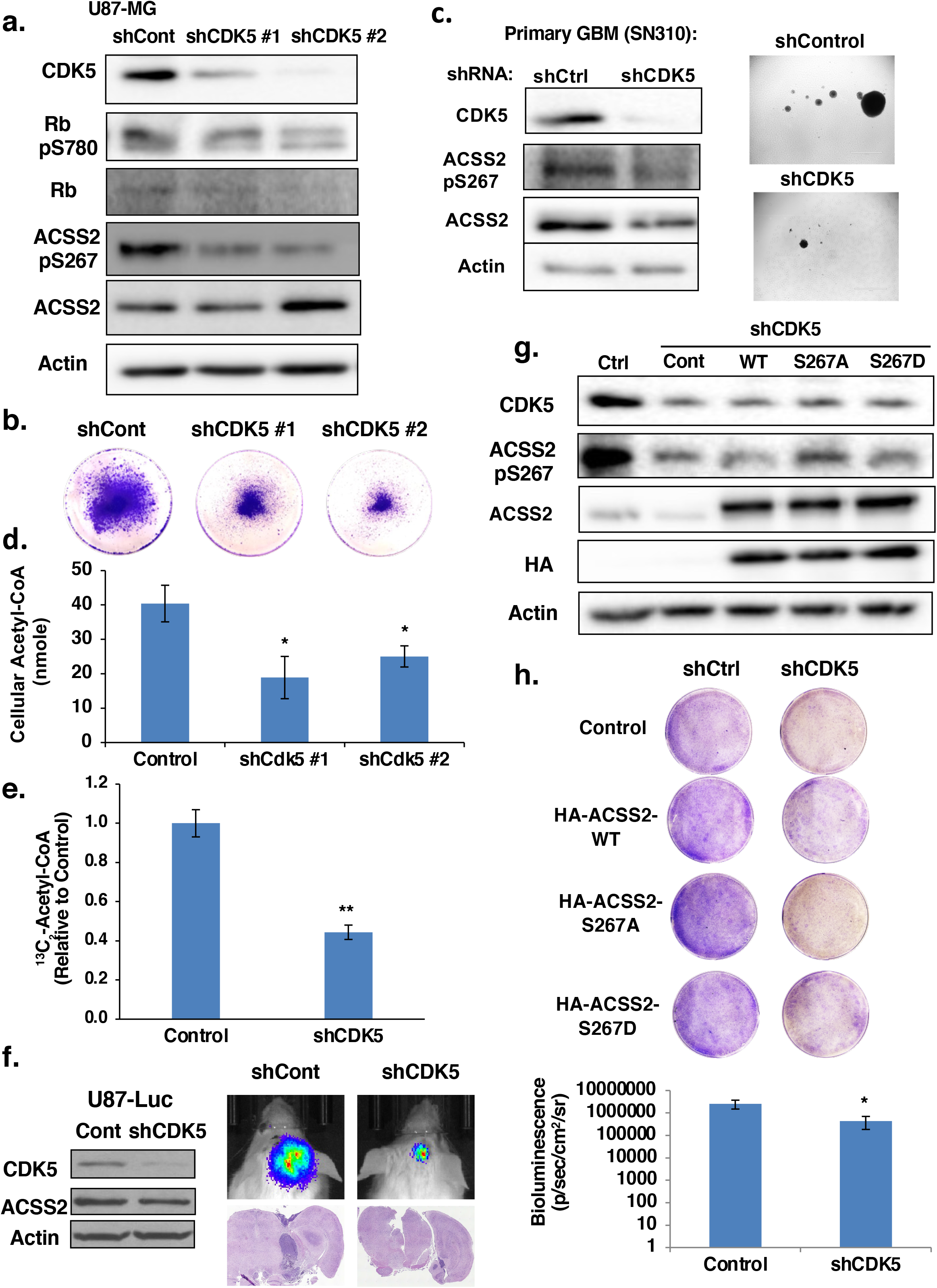
CDK5 is a critical regulator of GBM growth and requires ACSS2 phosphorylation. **(a)** Cell lysates from of U87-MG cells stably expressing shRNA against control or CDK5 were collected for immunoblot analysis with the indicated antibodies. **(b)** Representative images of U87-MG cells stained with crystal violet stably expressing of shRNA targeting control or CDK5. **(c)** Cell lysates from SN310 primary GBM cells expressing control or CDK5 shRNA were collected for immunoblot analysis with the indicated antibodies (left). Representative images from neurosphere growth assay (right) at day 6 comparing the neurosphere formation of control or CDK5 shRNA expressing SN310 cells. **(d)** Measurement of acetyl-CoA extracted from U87-MG cells stably overexpressing control or CDK5 shRNA. Student’s t-test reported as mean ± SEM. * = p-value < 0.05. **(e)** U87-MG cells stably expressing shRNA against control or CDK5 were labeled with ^13^C2-acetate for four hours and ^13^C2-acetyl-CoA concentrations were measured and shown. Student’s t-test reported as mean ± SEM. **p-value < 0.0001 **(f)** Cell lysate from U87-MG-Luciferase cells stably expressing control or CDK5 shRNA were collected for immunoblot analysis with the indicated antibodies (left). Representative images of tumor growth detected via bioluminescence and H&E staining at Day 16 following injection of U87-MG-luciferase cells expressing control or CDK5 shRNA (middle). Quantification of tumor size at Day 16 (shControl n=5, shCDK5 n=5) (right). Student’s t-test reported as mean ± SEM. * = p-value < 0.05. **(g)** Cell lysates from U87-MG cells stably expressing shRNA against control or CDK5 and overexpressing HA-ACSS2-WT (wildtype), HA-ACSS2-S267A, or HA-ACSS2-S267D mutants were collected for immunoblot analysis with the indicated antibodies. **(h)** Representative images of U87-MG cells stained with crystal violet containing shRNA targeting control or CDK5 and overexpressing ACSS2-WT (wildtype), ACSS2-S267A, or ACSS2-S267D mutants.

### OGT regulates acetate metabolism and GBM growth via ACSS2 S267 *in vitro* and *in vivo*

We have shown that GBM cell growth *in vitro* (Fig. 5e) and *in vivo* (Fig. 5f) requires ACSS2 Ser-267 phosphorylation and that CDK5-mediated regulation of cell growth also requires ACSS2 Ser-267 phosphorylation (Fig. 6h). To test whether OGT regulation of GBM lipid accumulation and growth also requires ACSS2 Ser-267 phosphorylation, we tested whether overexpression of wildtype ACSS2 or ACSS2 phospho-mimetic mutant would rescue the growth defects observed in OGT knockdown cells. Stable OGT knockdown reduced anchorage-independent growth *in vitro* could be rescued by overexpressing wildtype ACSS2 and ACSS2 S267D mutant, but not ACSS2 S267A mutant (Fig. 7a-7c). In addition, we found that overexpressing wildtype ACSS2 and ACSS2 S267D mutant, but not ACSS2 S267A mutant, could partly rescue nile red staining (Extended Data Fig. 7a). Consistent with the idea that OGT regulates acetyl-CoA via ACSS2, the OGT RNAi-mediated decrease in acetyl-CoA was reversed by overexpressing the ACSS2-267D mutant (Extended Data Fig. 7b, 7c). Similar to results *in vitro*, U87-MG cells expressing the phospho-mimetic mutant ACSS2 S267D, but not cells containing an ASCC2 S267A mutant, were able to partly rescue the growth effects of OGT depletion *in vivo* (Fig. 7d, 7e and Extended Data Fig. 7d). Thus, OGT regulation of GBM cell growth *in vivo* is partly dependent on ACSS2 Ser-267 phosphorylation.

**Fig 7.**
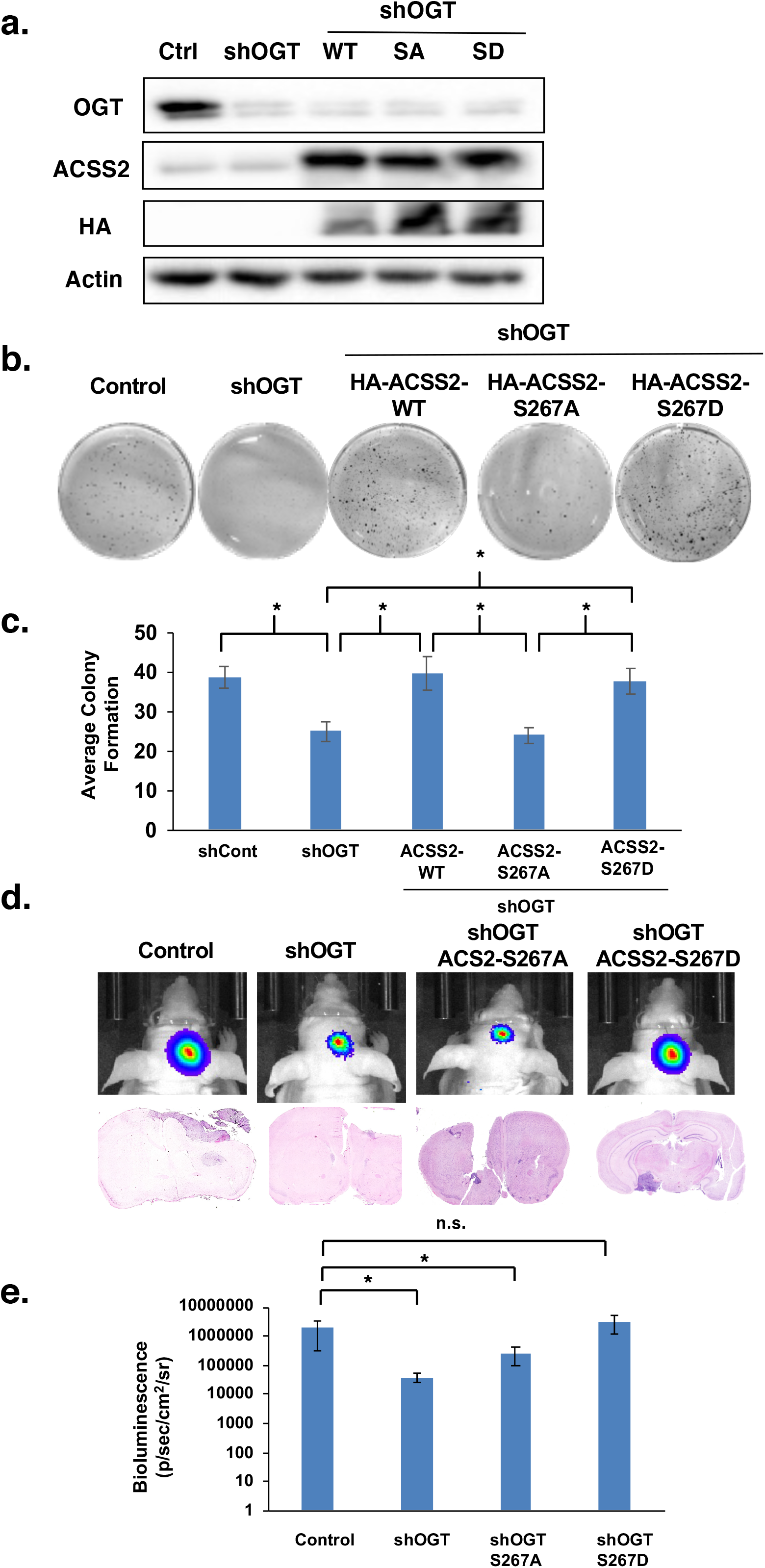
Phosphorylation of S267-ACSS2 is required for OGT-mediated GBM growth. **(a)** Cell lysates from of U87-MG cells stably expressing shRNA against control or OGT and overexpressing HA-ACSS2-WT, HA-ACSS2-S267A, and HA-ACSS2-S267D were collected for immunoblot analysis with the indicated antibodies **(b)** Representative images of cells generated in (a) and seeded into anchorage-independent growth assay and imaged at day 14. **(c)** Average colony formation quantified and presented as average from three independent experiments corresponding to Figure 7b showing U87-MG cells stable expressing ACSS2-WT, ACSS2-S267A and ACSS2-S267D mutants and expressing shControl or shOGT as indicated. Student’s t-test reported as mean ± SEM. * = p-value < 0.05. **(d)** Representative images of tumor growth detected via bioluminescence (top) and H&E staining (bottom) U87-MG-luciferase cells subjected to the same treatment as in (a) and then used for orthotopic intracranial injections in mice. **(e)** Quantification of tumor size at Day 16 corresponding to Figure 7d (shControl n=5, shOGT n=8, shOGT+ACSS2-S267A n=4, shOGT+ACSS2-S267D n=4). Student’s t-test reported as mean ± SEM. * = p-value < 0.05.

### Pharmacologically targeting OGT and CDKs in GBM *ex vivo*

To test whether targeting OGT or CDK5 in a preformed GBM tumor can block cancer growth, we treated *ex vivo* brain slices containing GBM tumors with an OGT inhibitor or pan-CDK inhibitor. Twelve days following intracranial injections of U87-MG-luc cells, intact brains were excised and sectioned, then slices containing tumors were cultured and treated with vehicle or Ac-5S-GlcNAc. Tumors in brain slices treated with control continued growing *ex vivo*, while tumors exposed to OGT inhibitor treatment significantly decreased in size (Fig. 8a, 8b) and exhibited reduced detection of the proliferation marker Ki-67, ACSS2 Ser-267 staining, and increased caspase-3 staining (Fig. 8a).

**Fig 8.**
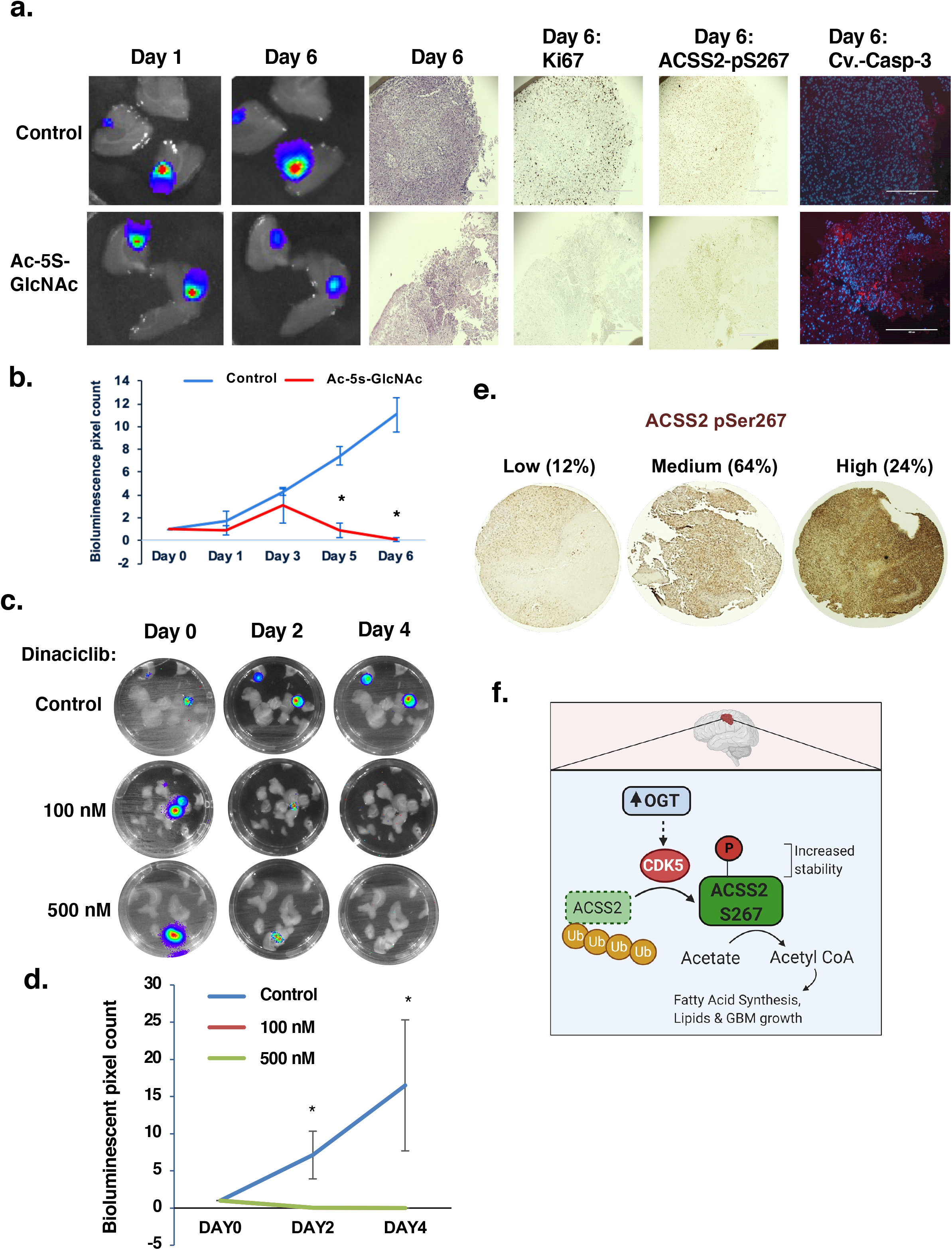
Targeting OGT or CDKs blocks GBM growth *ex vivo*. **(a)** Representative images depicting tumor growth in organotypic brain slices derived from mice intracranially injected with U87-MG-luc cells detected via bioluminescence. Brain slices containing tumors are treated with vehicle Control (DMSO) or Ac-5S-GlcNAc (200 μM) for indicated days (top) (image magnification 10x, scale bar: 400 μm). Slices were fixed and assayed for H&E, Ki-67, phospho-ACSS2-S267, cleaved caspase-3 staining. **(b)** Quantification of tumor growth at indicated day (Control: DMSO n=3, Ac-5S-GlcNAc n=3) (bottom). Student’s t-test reported as mean ± SEM. * = p-value < 0.05. **(c)** Luciferase images of *ex vivo* brain explants containing preformed tumors U87-MG-luc treated with indicated dose of Dinaciclib for indicated days. **(d)** Quantification of tumor growth at indicated day treated with Control: DMSO (n=3), dinaciclib 100 nM (n=3) or 500 nM (n=3). Student’s t-test reported as mean ± SEM. * = p-value < 0.05. **(e)** Immunohistochemical staining for ACSS2 pSer267 on a tissue microarray of Grade IV glioblastoma patient biopsies (n=69). **(f)** Model. Increased levels and activity of OGT in GBM leads to a CDK5-depepndent phosphorylation of ACSS2 on Ser267 that increases its stability and blocks ubiquitination and increases acetate conversion to acetyl-CoA and contributes to growth and survival of GBM tumors.

Importantly, treating control brain slices with the OGT inhibitor reduced total O-GlcNAc levels (Extended Data Fig. 7e) but did not alter brain tissue viability compared to control (Extended Data Fig. 7d). Treatment of GBM primary cells SN310 with pan-CDK inhibitor dinaciclib, which targets CDK1, CDK2, CDK5 and CDK9 (Parry et al., 2010a), blocked ACSS2 S267 phosphorylation (Extended Data Fig. 8a) and reduced neurosphere formation (Extended Data Fig. 8b). Treatment of U87-MG with dinaciclib also reduced ACSS2 S267 phosphorylation (Extended Data Fig. 8c) and also significantly decreased size of preformed tumors *ex vivo* (Fig. 8c, 8d) without causing loss of viability in brain slices (Extended Data Fig. 8d). These results demonstrate that targeting OGT and CDKs is efficacious in reducing preformed GBM tumor growth *ex vivo* and that this reduction is associated with reduced ACSS2 Ser-267 phosphorylation.

Lastly, we performed IHC analysis on glioblastoma (Grade IV) patient samples and found that phosphorylation levels of ACSS2 S267 was highly elevated in 24% percent of glioblastoma samples (Fig. 8e). These results support the idea that ACSS2 S267 phosphorylation is highly elevated in glioblastoma patients.

## Discussion

Our studies reveal a novel role of O-GlcNAc in regulation of acetate conversion to acetyl-CoA in glioblastoma cells via regulation of ACSS2. The brain has a unique ability to rewire its metabolism in response to changing metabolite availability (Magistretti and Allaman, 2015). Similarly, brain tumors must also be able to adapt in their ability to generate energy from non-glucose sources including acetate (Schild et al., 2018). In the present study, we demonstrated that elevated OGT and O-GlcNAcylation helps glioblastomas rewire metabolism to utilize acetate that provides a survival and growth advantage in this unique environment. Specifically, we propose a previously unknown pathway by which OGT regulates phosphorylation of ACSS2 on serine 267 in a CDK5-dependent manner and that this phosphorylation stabilizes ACSS2 protein levels, reduces its polyubiquitination, and regulates acetate conversion into acetyl-CoA and contributes to GBM lipid storage and growth (Fig. 8f). To our knowledge, this is the first report to link CDK5 directly to regulation of cancer cell metabolism.

Interestingly, CDK5 has previously been shown to be regulated by glucose (Lowman et al., 2010; Ubeda et al., 2004). In pancreatic beta-cells, increased concentrations of glucose result in increased CDK5 activity that regulates insulin gene expression (Ubeda et al., 2004). However, the mechanism by which glucose regulates activation of CDK5 is not known. We speculate that increased glucose may lead to elevated O-GlcNAcylation and activation of CDK5 activity. Intriguingly, three residues on CDK5 have been shown to be dynamically O-GlcNAc modified, directly regulating its ability to activate p53-mediated apoptosis in neurons during intracerebral hemorrhage (Ning et al., 2017). It will be of interest to determine whether CDK5 is directly O-GlcNAcylated in glioblastomas or whether O-GlcNAc regulates CDK5 via an indirect mechanism. Despite OGT inhibitors not being well developed to determine tumor efficacy or possible toxicities, pan-CDK inhibitors such as dinaciclib have shown promise as a primary therapy for multiple myeloma (Kumar et al., 2015) and in preclinical models of pancreatic (Feldmann et al., 2011) and ovarian cancers (Chen et al., 2015). Therefore, in light of our *ex vivo* results which suggest inhibition of glioblastoma growth by dinaciclib, it would be of clinical interest to analyze the efficacy of these compounds, as well as more CDK5-specific inhibitors, for targeting glioblastoma metabolism and growth.

Acetate conversion to acetyl-CoA in cancer can contribute to three major metabolic pathways: *de novo* fatty acid and isoprenoid synthesis, the TCA cycle and histone acetylation (Schug et al., 2016). For example, nuclear ACSS2 has been described as having the ability to recapture acetate produced as the result of histone deacetylation and can utilize this acetate to maintain histone acetylation (Bulusu et al., 2017). In addition, acetyl-CoA locally produced by ACSS2 using acetate generated from nuclear protein deacetylation can be used for acetylation of promoters critical for regulating gene expression of genes involved in cancer cell survival (Li et al., 2017). It would be intriguing to explore the possibility that OGT-CDK5-mediated Ser267 phosphorylation may also regulate the localization of ACSS2, influencing gene expression processes through ACSS2-dependent histone acetylation (Bulusu et al., 2017; Li et al., 2017). Our data does suggest that the OGT-CDK5-ACSS2 pathway may contribute to lipid biosynthesis as OGT and CDK5 knockdown inhibition of intracellular lipid storage can be reversed by overexpressing ACSS2 phospho-mimetic mutant. Recent studies have shown that cancers in the brain must adapt to lack of lipid availability in the brain environment (Jin et al., 2020) and thus are highly dependent on fatty acid synthesis for growth and survival (Ferraro et al., 2021). Our study provides one potential pathway for a mechanism by which GBM cells can generate acetyl-CoA, a major metabolite feeding lipid synthesis, via phosphorylation of ACSS2 by CDK5 and activation of nutrient sensor OGT. Future experiments investigating how OGT-CDK5-ACSS2 contributes to acetate metabolism-dependent regulation of fatty acid biosynthesis, the TCA cycle and histone acetylation will be further explored.

## Methods

### Cell Culture

U87-MG and T98G cell lines were obtained from ATCC (Manassas, VA, USA) and cultured according to ATCC protocol supplemented with 10% fetal bovine serum (FBS), 5% 10,000 Units/mL Penicillin-10,000 μg/mL Streptomycin, and 5% 200 mM L-Glutamine. Primary GBM WHO grade IV cell lines SN186 (76 yr old male), SN275 (58 yr old male) and SN310 (78 yr old female) were provided as a kind gift by P. Hothi (Swedish Neuroscience Institute, Seattle, WA) and have been previously described (Hothi et al., 2012). Primary GBM cell lines were cultured in Neurobasal-A Medium (Gibco; Gaithersburg, MD) supplemented with 1x B27 Supplement (Gibco), 1 mM sodium pyruvate, 0.5 mM L-glutamine, 1x Antibiotic-Antimycotic (Gibco), 0.02 mg/mL insulin solution, 20 ng/mL EGF, and 20 ng/mL FGF-2. For crystal violet staining, 5 x 10^4^ cells were plated and subjected to the treatments as described in the individual figures and then stained with 0.5% crystal violet prepared in a 1:1 methanol-water solution followed by PBS washes.

### Animal models of cancer

Nu/Nu athymic 6-8 week old mice from Charles River Laboratories (Wilmington, MA, USA) were immobilized using the Just for Mice ™ Stereotaxic Frame (Harvard Apparatus, Holliston, MA, USA) and injected intracranially with 5 μl of a 20,000 U87-MG-Luciferase cells/μl solution. Tumor growth was monitored via bioluminescence imaging on the IVIS 200 system (Perkin Elmer) and results analyzed using Living Image software (Caliper Life Sciences, Waltham, MA, USA). For survival studies, mice were injected and health monitored until in moribund state. All animal protocols were approved by the Institutional Animal Care and Use Committee.

### Human samples

GBM tumor tissue samples were obtained from 36 patients diagnosed with craniotomy resection of the tumor. All patients received standard 6-week radiation treatment with concurrent temozolomide. Specimens from multiple resections were included in TMA including 26 samples from 36 patients that were isolated in the first resection, 30 samples from same 36 patients from second resections, and 13 samples from the 36 patients were isolated from third resections thus total of 69 samples were analyzed. Histology was evaluated according to the World Health Organization (WHO) classification for CNS tumors. For survival analysis Kaplan–Meier curves were generated using the online database UALCAN (http://ualcan.path.uab.edu/analysis.html) (Chandrashekar et al., 2017)to determine the relevance of CDK5 mRNA expression in patients with GBM.

### Reagents

Anti-actin (Santa Cruz Biotechnology; Dallas, TX, USA), anti-OGT, anti-ACSS2, anti-Cdk5, anti-K48 polyubiquitin, anti-HA-tag, anti-p35/p25, anti-Rb, anti-phospho Rb-S780, anti-cleaved-caspase-3, IgG-conjugated sepharose beads (Cell Signaling Technology; Danvers, MA, USA), anti-O-GlcNAc (Sigma Aldrich; St. Louis, MO, USA) and anti-thiophosphate ester (Abcam; Cambridge, U.K.). Protein G sepharose 4B beads (Invitrogen; Carlsbad, CA, USA). Plasmids used were pReceiver N-Flag OGT, pReceiver HA-ACSS2, HA-ACSS2-S267A, HA-ACSS2-S267D (Genecopoeia; Rockville, MD, USA). Cdk5-HA was a gift from Sander van den Heuvel (Addgene plasmid # 1872). HA-Ub and HA-Ub-K48R was a gift from Edward Hartsough (Drexel University). Ac-5S-GlcNAc and NButGT were a kind gift from David J. Vocadlo and previously described (Gloster et al., 2011). Cycloheximide (Sigma Aldrich, St. Louis, MO, USA), Dinaciclib (Selleckchem, Houston, TX, USA), lactacystin (Calbiochem, San Diego, CA, USA), D-luciferin potassium salt (Perkin Elmer, Waltham, MA, USA), and crystal violet (Sigma Aldrich, St. Louis, MO, USA). pReceiver.HA-ACSS2-WT, S267A, and S267D plasmid were provided by Genewiz (South Plainfield, NJ, USA) and confirmed via Sanger sequencing.

Rabbit polyclonal antibody recognizing phosphorylated ACSS2 S267 was created by YenZym Antibodies (South San Francisco, CA, USA). A peptide containing ACSS2 pS267 was injected into rabbits and serum was collected and purified using affinity column conjugated with nonphosphorylated ACSS2 S267 peptide to exclude antibodies recognizing ACSS2 nonphosphorylated form, followed by an affinity column conjugated with phosphorylated ACSS2 S267 peptide to purify the ACSS2 S267 phospho-specific antibody. This antibody was the eluted and used at concentration of 0.22 μg/ml.

### RNA interference

Stable cell lines for shRNA knockdowns were generated as previously described (Caldwell et al., 2010). Briefly, HEK-293T cells were grown to ∼75% confluency and transfected. Prior to transfection, 20 μg of shRNA or overexpression plasmid DNA, 10 μg VSVG, 5 μg RSV, and 5 μg RRE were gently mixed and incubated in 1.5 mL of Optimem for 5 minutes. Concurrently, 105 μl of PEI was added dropwise to 1.5 mL of Optimem and incubated for 5 minutes. Following the 5 minute incubation, the PEI solution was added dropwise to the DNA solution and incubated for at least 30 minutes. The PEI-DNA solution was then added dropwise to the HEK-293T cells already plated with 5 mL of Optimem and the cells were incubated overnight in the transfection media. Approximately 16-18 hours later, the transfection media was replaced with normal growth media. Viral supernatants were collected at 24 and 48 hours following the media change. These supernatants were passed through a 0.45 μm filter and portioned into 1 mL aliquots to be stored at −80°C if not for immediate us.

Control shRNA was acquired from Addgene (plasmid 1864), from D. Sabatini (Massachusetts Institute of Technology). Control-scrambled shRNA sequence used was: CCTAAGGTTAAGTCGCCCTCGCTCTAGCGAGGGCGACTTAACCTT. All other shRNA constructs were acquired from Sigma and shRNA sequence used: for OGT-1, GCCCTAAGTTTGAGTCCAAATCTCGAGATTTGGACTCAAACTTAGGGC, and for OGT-2, GCTGAGCAGTATTCCGAGAAACTCGAGTTTCTCGGAATACTGCTCAGC. Cdk5 shRNA sequence used: for Cdk5-1, CCGGGTGAACGTCGTGCCCAAACTCCTCGAGGAGTTTGGGCACGACGTTCACTTTTTTG, and for Cdk5-2, CCGGCAGAACCTTCTGAAGTGTAACCTCGAGGTTACACTTCAGAAGGTTCTGTTTTTTG. ACSS2 shRNA sequence used: CCGGGCTTCTGTTCTGGGTCTGAATCTCGAGATTCAGACCCAGAACAGAAGCTTTTTG.

### mRNA expression

RNA was isolated from cells using RNEasy products and protocols (Qiagen, Valencia, CA, USA). Taqman gene expression assay primer probes for OGT (Hs00269228_m1), ACSS2 (Hs00218766_m1), and GAPDH (Hs02758991_g1) were purchased from Applied Biosystems (Foster City, CA, USA). qRT-PCR was performed as previously described (Sodi et al., 2015) using Brilliant II qRT-PCR Master Mix 2 Kit (Stratagene, San Diego, CA, USA) using Applied Biosystems 7500 machine.. Briefly, isolated RNA concentrations were determined by measuring on a NanoDrop and all samples were diluted to match the sample with the lowest concentration. A mastermix was then generated for the individual primers using 8.8 μl dH2O, 12.5 μl Brilliant PCR master mix, 0.4 μl diluted dye (0.5 μl dye into 250 μl dH2O), 0.1 μl reverse transcriptase, and 1.25 μl primer per sample. A total reaction volume of 25 μl was used by gently mixing 2 μl of sample RNA and 23 μl of the mastermix into the reaction tubes, being careful to avoid bubbles. The samples were then loaded into the Applied Biosystems 7500 machine and the experimental setup was as follows: acquire experiment as “Quantitation ΔΔCT”, targets and samples were assigned, the initial holding stage adjusted to 50°C for 30 minutes, followed by a secondary holding stage at 95°C for 10 minutes, then 40 cycles consisting of 95°C for 15 seconds and cooling to 60°C for 1 minute were completed. Data were exported from the Applied Biosystems software and expression levels were analyzed using Data Assist v2.0 (Life Technologies, Grand Island, NY, USA).

### Immunoblotting

Immunoblotting protocols have been previously described (Lynch et al., 2012). Briefly, cell lysates from 1-5 x 10^6^ cells were prepared in radioimmune precipitation assay buffer (150 mM NaCl, 1% NP40, 0.5% DOC, 50 mM Tris HCL at pH 8, 0.1% SDS, 10% glycerol, 5 mM EDTA, 20 mM NaF, and 1 mM Na_3_VO_4_) supplemented with 1 μg/ml each of pepstatin, leupeptin, aprotinin, and 200 μg/ml PMSF. Lysates were cleared by centrifugation at 16,000 x *g* for 15 minutes at 4 °C and analyzed by SDS-PAGE and autoradiography with chemiluminescence. Proteins were analyzed by immunoblotting using primary antibodies indicated above.

### Immunohistochemical staining

The slides containing glioblastoma tumors were deparaffinized by Xylene and subsequent rehydration was done by decreasing concentrations of ethanol-water mixture. Antigen retrieval was done by citrate buffer immersion and steaming the slides for 45 minutes. Tissue was treated with 200-400 ul 1% BSA+5% serum PBS solution for 1hr. Primary incubation was done at 4° C overnight using OGT, O-GlcNAc RL2 (Santa Cruz 1:50), p-S267 ACSS2 (1:100) antibodies.

Secondary antibody incubation was done for an hour at RT with respective antibodies at 1: 200 dilution. The stain was developed using the DAB kit by Vectastain (Vector Labs, Burlingame, CA, USA). Finally, slides were mounted and imaged under a light microscope. The staining patterns of tumor tissues of GBM were assessed by a board-certified pathologist.

### Immunoprecipitation

Cells were lysed and extracted with radioimmune precipitation assay buffer (150 mM NaCl, 1% NP40, 0.5% DOC, 50 mM Tris HCL at pH 8, 0.1% SDS, 10% glycerol, 5 mM EDTA, 20 mM NaF, and 1 mM Na3VO4) supplemented with 1 μg/ml each of pepstatin, leupeptin, aprotinin, and 200 μg/ml PMSF. For each sample, 1 mg of protein lysate was added to an Eppendorf tube and precleared with 15 μl of prewashed protein G sepharose beads by rotating at 4°C for 30 minutes followed by a 1-minute centrifugation at 2,500 x g at 4°C. These precleared supernatants were transferred to fresh tubes and were incubated with the indicated antibodies and lysates overnight at 4^0^C. The following day, 30 μl of prewashed protein G sepharose beads were added to the immunoprecipitations and incubated for 2-3 hours by rotating at 4°C. The immunoprecipitation reactions were centrifuged at 2,500 x g at 4°C for 5 minutes followed by a wash with 750 μl of 1X PBST and this process was repeated three more times. After the final wash, the antibody-protein-bound beads were collected via centrifugation at 4°C and removal of supernatant. Beads were resuspended in 5x SDS loading buffer, boiled for 5 min and loaded. Immunoprecipitated proteins were resolved by SDS-PAGE sent for mass spectrometry or transferred to a PVDF membrane. Immunoblotting protocol was followed as per methods above.

### LC-MS/MS Analyses and Data Processing

Quantitative comparisons of post-translational modifications via mass spectrometric analysis on immunoprecipitated HA-ACSS2 treated with DMSO or 100 μM NButGT was performed by The Wistar Institute Proteomics & Metabolomics facility (Philadelphia, PA, USA). Briefly, liquid chromatography tandem mass spectrometry (LC-MS/MS) analysis was performed using a Q Exactive Plus mass spectrometer (ThermoFisher Scientific, Waltham, MA, USA) coupled with a Nano-ACQUITY UPLC system (Waters). Samples were digested in-gel with trypsin and injected onto a UPLC Symmetry trap column. Tryptic peptides were separated by reversed phase HPLC on a BEH analytical column. Eluted peptides were analyzed by the mass spectrometer.

Peptide sequences were identified using MaxQuant 1.5.2.8 (Cox and Mann, 2008). MS/MS spectra were searched against a UniProt human protein database. Consensus identification lists were generated with false discovery rates of 1% at protein, and peptide levels.

### Soft-Agar Colony Formation Assay and Neurosphere Assay

Soft agar assays have been previously described (Lynch et al., 2012). Briefly, cells (1 x 10^4^/well) infected with lentiviral shRNA or overexpression vectors, selected and plated in triplicate on top layers consisting of growth medium containing 0.3% agarose. The top layer was overlaid with 1 ml of medium. The cells were grown for 14 days, having the media changed every other day. After 14 days, the colonies were stained with 1 ml of 0.05% p-iodonitrotetrazolium violet overnight, and colonies measuring > 50 μm were counted manually. For neurosphere assays, SN310 primary glioblastoma cells (1 x 10^3^) infected with shRNA, selected and were plated in polyhema-coated (1.2%) 24-well plates with the number of neurospheres formed being determined at Day 14.

### In Vitro Kinase Assay

Wildtype ACSS2 and S267A mutant were cloned into a pET28a vector into the NotI and BamH1 sites and transformed into BL21 cells. The BL21 cultures were induced with 1mM IPTG at 37°C for 3 hours. Cells were centrifuged and resuspended in His Lysis buffer (50mM Tris, pH 8.0, 5 mM imidazole pH 8.0, 500mM NaCl), supplemented with protease inhibitors (1 μg/mL each of pepstatin, leupeptin, and aprotinin and 200 μg/mL phenylmethylsulfonylfluoride (PMSF) and 100mg/ml lysozyme. The lysate was sonicated on ice and centrifuged at 15000 x g for 15 minutes at 4°C. The supernatant was rotated over Nickel NTA Agarose beads (Gold Biotechnology, Cat# H-350-25) in a column (Gold Biotechnology, Cat# P-301) for 2 hours at 4°C. Upon collecting flow-through, beads were washed with wash buffer (50 mM Tris, pH 8.0, 500 mM NaCl, 40 mM imidazole, pH 8.0), and eluted with His Elution buffer (500mM NaCl, 50mM Tris, pH 8.0, 250 mM imidazole, pH 8.0). Dialysis was performed in Slide-A-Lyzer G2 Dialysis Cassette (Thermo Scientific, Prod# 88251) for 2 hours at 4°C in dialysis buffer (20 mM Tris, pH 8.0, 150 mM NaCl, 1 mM DTT). Recombinant CDK5/p25 (60 ng) (EMD Millipore, Burlington, MA, USA) was combined with purified ACSS2 (2ug) in kinase buffer (50 mM HEPES, pH 7.5, 0.625 mM MgCl_2_, 0.625 mM MnCl_2_, 12.5 mM NaCl) and 1mM ATPγS. The kinase reaction incubated for 5 minutes at 30°C, then was alkylated with 2.5mM PNBM. The alkylating reaction proceeded for 1 hour at room temperature, followed by the addition of 6× boiling sample buffer to stop the reaction. Reactions were then resolved by SDS–PAGE, followed by immunoblotting with the indicated antibodies.

### Nile Red Staining of Cells

Cells were fixed using 4% Formalin and washed in 1xPBS prior to staining. Working solution (5 μg/ml) in PBS was gently added to cells and incubated for 30 minutes away from light. Cells were then washed three times in 1x PBS and visualized and photographed on EVOS FL (Life technologies) using Texas Red filter.

### Free Fatty Acid and Acetyl-CoA Extraction and Quantification

Free Fatty Acid concentrations were measured with the Free Fatty Acid Quantification Kit (BioVision Inc., Milpitas, CA, USA) according to the manufacturer’s protocol. Acetyl-CoA concentrations were measured with the Acetyl-Coenzyme A Assay Kit (Sigma Aldrich) according to the manufacturer’s protocol.

### ^13^C_2_-Acetate Labeling

Cells were grown to ∼80% confluency in a 6 cm dish in normal growth medium. 24 hours prior to labeling cells were grown in serum-free DMEM. Cells were incubated for 4 hours in 100 μM 13C_2_-sodium acetate. Cells were collected in cold PBS then centrifuged at 1,000 rpm for 5 minutes at 4 °C. Acetyl-CoA was then extracted from the cell pellets using 80% methanol – 20% dH2O and the samples were then analyzed via LC/MS.

### *Ex vivo* brain slice model

Organotypic hippocampal cultures were prepared as described previously (Jackson et al., 2014) with some modifications. Briefly, adult mice (6-8 week) or mice after 12 days following intracranial GBM injection, as described above, were decapitated and their brains rapidly removed into ice-cold (4°C) sucrose-aCSF composed of the following (in mM): 280 sucrose, 5 KCl, 2 MgCl2, 1 CaCl2, 20 glucose, 10 HEPES, 5 Na^+^-ascorbate, 3 thiourea, 2 Na^+^-pyruvate; pH=7.3. Brains were blocked with a sharp scalpel and sliced into 250 µm sections using a McIlwain-type tissue chopper (Vibrotome inc). Four to six slices were placed onto each 0.4 µm Millicell tissue culture insert (Millipore) in six-well plates, 1 ml of medium containing the following: Neurobasal medium A (Gibco), 2% Gem21-Neuroplex supplement (Gemini), 1% N2 supplement (Gibco), 1% glutamine (Invitrogen), 0.5% glucose, 10 U/ml penicillin, and 100 μg/ml streptomycin (Invitrogen), placed underneath each insert. One-third to one half of the media was changed every 2 d. Tumor growth was monitored via bioluminescence imaging on the IVIS 200 system (Perkin Elmer) and results analyzed using Living Image software (Caliper Life Sciences, Waltham, MA, USA). For MTS assay, individual brain slices were transferred to a 96-well plate and subjected to Promega CellTiter 96® Aqueous One Solution (Cat: G3582) mixed in a 1:5 ratio with culture media and treated as previously described (Mewes et al., 2012). Tissues were incubated at 37°, 5% CO_2_ for 4 hours and absorbance at 490nm was measured with Tecan Spark Microplate reader.

### Statistical Analysis and reproducibility

All results shown are results of at least three independent experiments and shown as averages and presented as mean ± s.e. P-values were calculated using a Student’s two-tailed test (* represents p-value ≤ 0.05 or **p-value ≤ 0.01 or as marked in figure legend). Statistical analysis of growth rate of mice was performed using ANCOVA. *p-value < 0.05.

## SUPPLEMENTAL INFORMATION

Supplemental information can be found online at….

## ACKNOWLEDGMENTS

This work was supported by PA CURE grant (to M.J.R., N.W.S and J.G.J.), Drexel University Dean’s Fellowship award (to Z.A.B. and L.C.). The authors thank Chaitali Bhadiadra for technical assistance, Dr. Valerie Sodi and Dr. Edward Hartsough for helpful discussions. In memoriam Christos D. Katsetos.

## Author contributions

L.C. and Z.A.B. performed most of the experimental work; J.J., R.A.M., and R.H.L. helped with experimental work; C.M.F helped establish intracranial mouse model; S.F., M.T.D., and N.W.S. performed acetate labeling and analyzed data. W.A.G., L.D. and C.D.K. helped with IHC and C.D.K. performed pathological analysis; W.S. provided GBM tissue array; J.G.J., L.C., R.A.M. helped establish *ex vivo* brain slice model. L.C., Z.A.B. and M.J.R participated in study conception and design as well as data analysis and interpretation; L.C., Z.A.B. and M.J.R drafted the manuscript; All co-authors reviewed the final manuscript version.

