## Supplemental Data for "O-GlcNAc transferase regulates glioblastoma acetate metabolism via regulation of CDK5-dependent ACSS2 phosphorylation"

### Supplementary Figure Legends

**Extended Figure 1. Inhibition of OGT or O-GlcNAcylation impedes glioblastoma growth. (a)** Representative images from anchorage-independent growth assay comparing the colony formation of control or OGT shRNA expressing U87-MG (top) and T98G (bottom) cells. **(b)** U87-MG cells were treated with 100 μM and 200 μM of OGT inhibitor, Ac-5S-GlcNAc, for 24 hours and then harvested for lysate (top) or stained with crystal violet (bottom). **(c)** Cell lysate from U87-MG cells expressing luciferase and control or OGT shRNA were collected for immunoblot analysis with the indicated antibodies. **(d)** Control or stably overexpressing OGT U87-MG cells were placed in soft agar assay. Representative images from anchorage-independent growth assay.

**Extended Figure 2. Targeting OGT expression inhibits lipid metabolism. (a)** Cell lysates from of U87-MG cells stably expressing control shRNA or shOGT were collected for immunoblot analysis with the indicated antibodies. **(b)** Representative images of nile red staining of U87-MG cells under same conditions as in (a). **(c)** Measurement of relative free fatty acids in U87-MG cells stably overexpressing shControl or shOGT. Student's t-test reported as mean ± SEM. * = p-value < 0.05. **(d)** Measurement of acetyl-CoA extracted from U87-MG cells stably overexpressing shControl or shOGT. Student's t-test reported as mean ± SEM. * = p-value < 0.05. **(e)** Cell lysates from T98G cells stably expressing control shRNA or shOGT were collected for immunoblot analysis with the indicated antibodies. **(f)** Representative images of nile red staining of T98G cells under same conditions as in (e). **(g)** Measurement of relative free fatty acids in T98G cells stably overexpressing shControl or shOGT. **(h)** Measurement of acetyl-CoA extracted from T98G cells stably overexpressing shControl or shOGT. **(i)** Cell lysate from T98G cells that were treated with control (DMSO) or 100 μM OGA inhibitor (NButGT) for 48 hours were collected for immunoblot analysis with the indicated antibodies (top) and measurement of acetyl-CoA extracted from the treated T98G cells is shown (bottom). Student's t-test reported as mean ± SEM. * = p-value < 0.05

**Extended Figure 3. Elevated OGT/O-GlcNAcylation increases ACSS2 protein levels. (a)** Total RNA was collected from U87-MG cells stably expressing shControl or shOGT. Quantification of qRT-PCR performed on RNA extracts analyzing OGT and ACSS2 gene expression normalized to GAPDH. Student's t-test reported as mean ± SEM. * = p-value < 0.05. **(b)** Graphical representation of densitometry from three independent experiments comparing ACSS2 protein expression normalized to actin in lysate from control and OGT overexpressing U87-MG cells as shown in Fig. 3a. Student's t-test reported as mean ± SEM. * = p-value < 0.05. **(c)** Cell lysates from SN310 cells overexpressing control or OGT were collected for immunoblot analysis with indicated antibodies. **(d)** Cell lysates from U87-MG cells treated with OGA inhibitor (NButGT) for indicated times and dose were collected for immunoblot analysis with the indicated antibodies. **(e)** Cell lysate collected from U87-MG cells stably overexpressing control or HA-ACSS2 and treated with either DMSO or 100 μM NButGT for 48 hours. The lysates were then subjected to immunoprecipitation with isotype control IgG or HA-tag antibodies and analyzed via SDS-PAGE for further analysis via western blotting and mass-spectrometry. **(f)** Scansite analysis of the ACSS2 protein sequence. (**g)** Ser-267 and the consensus motif (underlined) are evolutionarily conserved on ACSS2 protein in mammals. **(h)** Non-radioactive *in vitro* kinase assay performed with increasing concentrations of recombinant ACSS2 with recombinant CDK5. **(i)** Non-radioactive *in vitro* kinase assay performed with either wild-type or S267A purified ACSS2 in presence or absence of recombinant CDK5. **(j)** Immunoprecipitation was performed with indicated antibodies from U87-MG cells stably expressing wild-type (WT) or S267A HA-ACSS2. **(k)** *In vitro* immunoprecipitation of recombinant CDK5 in presence of purified ACSS2 and immunoblot analysis with the indicated antibodies. **(l)** Immunoprecipitation of endogenous CDK5 from U87-MG glioblastoma cells treated with 100 μM NButGT for 48 hours and immunoblot analysis with the indicated antibodies.

**Extended Figure 4. ACSS2-serine 267 phosphorylation in GBM cells and neuronal tissue. (a)** Cell lysates from U87-MG cells were subjected to immunoprecipitation with isotype control IgG or phospho-ACSS2-S267 antibody and analyzed via SDS-PAGE for further analysis via western blotting with indicated antibodies. **(b)** U87-MG cells were treated with DMSO, 100 μM or 200 μM of OGT inhibitor, Ac-5S-GlcNAc, for 48 hours and then lysate were collected for immunoblot analysis with the indicated antibodies. **(c)** Cell lysates from normal human astrocytes, T98G or U87-MG cells treated with control (DMSO) or OGA inhibitor (NButGT) (100 μM, 48 hrs) were collected for immunoblot analysis with the indicated antibodies. **(d)** Cell lysates from *ex vivo* brain slice from P10 mice were treated with control (DMSO) or OGA inhibitor (NButGT) (100 μM, 48 hrs) were collected for immunoblot analysis with the indicated antibodies. **(e)** Graphical representation of densitometry from three independent experiments comparing ratio of K48 polyubiquitination to HA from immunoprecipitation experiments of WT-, S267A-, and S267D-ACSS2 shown in Figure 5b. Student's t-test reported as mean ± SEM. * = p-value < 0.05, **= p-value < 0.01, ***= p-value < 0.005. **(f)** Cell lysate from U87-MG-Luciferase cells stably expressing shRNA against endogenous ACSS2 (left) and overexpressing HA-ACSS2-WT or HA-ACSS2-S267A (right) were collected for immunoblot analysis with the indicated antibodies.

**Extended Figure 5. CDK5 suppression inhibits GBM tumorigenic potential. (a)** Kaplan-Meier overall survival curve for GBM cancer patients according to CDK5 high (n=38) or low (n=114) expression levels. **(b)** Representative images from anchorage-independent growth assay (top) comparing the colony formation of control or CDK5 shRNA expressing U87-MG cells. Data are quantified and presented as average from three independent experiments (bottom). Student's t-test reported as mean ± SEM. * = p-value < 0.05. **(c)** Cell lysates from T98G cells expressing control or CDK5 shRNA where collected for immunoblot analysis with the indicated antibodies (top). Representative images from cells stained with crystal violet (bottom). **(d)** Cells were placed in anchorage-independent growth assay (top) comparing the colony formation of control or CDK5 shRNA expressing T98G cells. Data are quantified and presented as average from three independent experiments (bottom). Student's t-test reported as mean ± SEM. * = p-value < 0.05. **(e)** Neurospheres were quantified for SN310 cells expressing control or CDK5 shRNA. Student's t-test reported as mean ± SEM. * = p-value < 0.05. **(f)** U87-MG cells stable expressing control or shRNA targeting CDK5 were imaged following nile red staining.

**Extended Figure 6. CDK5 is required for OGT-mediated effect on GBM growth. (a)** Cell lysate from U87-MG cells stably overexpressing control or OGT and control or CDK5 shRNA were collected for immunoblot analysis with the indicated antibodies. **(b)** Representative images from anchorage-independent growth assay (top) comparing the colony formation of control and OGT overexpressing cells expressing control or CDK5 shRNA U87-MG cells. Data are quantified and presented as average from three independent experiments (bottom). Student's t-test reported as mean ± SEM. * = p-value < 0.05. **(c)** Cell lysate from U87-MG cells stably overexpressing control or OGT and control or CDK5 shRNA were collected for immunoblot analysis with the indicated antibodies. **(d)** Measurement of acetyl-CoA extracted from U87-MG cells stably overexpressing control or OGT and control or CDK5 shRNA. Student's t-test reported as mean ± SEM. * = p-value < 0.05. **(e)** Cell lysate from U87-MG cells that were treated with control (DMSO) or 100 µM OGA inhibitor (NButGT) for 48 hours stably expressing control or CDK5 shRNA were collected for immunoblot analysis with the indicated antibodies. **(f)** Measurement of acetyl-CoA extracted from cells treated with control (DMSO) or 100 µM OGA inhibitor (NButGT) for 48 hours and stably expressing control or CDK5 shRNA. **(g)** Representative images of nile red staining of U87-MG cells stably expressing ACSS2-WT, ACSS2-S267A and ACSS2-S267D mutants and expressing shCDK5 as shown in Fig. 6g.

**Extended Figure 7. ACSS2-S267D mutant rescues CDK5 and OGT-mediated inhibition of GBM growth. (a)** Representative images of nile red staining of U87-MG cells stable expressing ACSS2-WT, ACSS2-S267A and ACSS2-S267D mutants and overexpressing shControl or shOGT as shown in Figure 7a. **(b)** Cell lysates from U87-MG cells stably overexpressing control or ACSS2-S267D mutant and shRNA against control or OGT were collected for immunoblot analysis with the indicated antibodies. **(c)** Measurement of acetyl-CoA extracted from U87-MG cells stably overexpressing control or ACSS2-

S267D mutant and shRNA against control or OGT. Student's t-test reported as mean ± SEM. * = p-value < 0.05. **(d)** Cell lysate from U87-MG-luciferase cells stably expressing HA-ACSS2-S267A and HA-ACSS2-S267D and control or OGT shRNA were collected for immunoblot analysis with the indicated antibodies. (**e)** Cell lysates from *ex vivo* brain slice from adult mice were treated with control (DMSO) or OGT inhibitor (Ac-5S-GlcNAc) (200 μM, 48 hrs) were collected for immunoblot analysis with the indicated antibodies. (**f)** *Ex vivo* brain slice as treated in Fig. 8A (without tumor) were collected at day 6 and analyzed for cell viability (MTS assay). Student's t-test reported as mean ± SEM. * = p-value < 0.05.

**Extended Figure 8. Dinaciclib treatment of GBM cells blocks ACSS2-Ser267 phosphorylation, neurosphere formation, and tumor growth *ex vivo*. (a)** Lysates from SN310 primary GBM cells treated for 48 hours with indicated concentrations of dinaciclib or vehicle (DMSO) were collected for immunoblot analysis with the indicated antibodies. **(b)** Representative images from SN310 neurosphere formation (top) and quantification of neurosphere formation of primary SN310 GBM cells treated with dinaciclib in (a) (bottom). Student's t-test reported as mean ± SEM. * = p-value < 0.05. **(c)** Lysates from U87-MG cells treated for 48 hours with indicated concentrations of dinaciclib or vehicle (DMSO) were collected for immunoblot analysis with the indicated antibodies. **(d)** *Ex vivo* brain slice were treated with indicated dose of dinaciclib or vehicle (DMSO) as in Fig. 8c (without tumor) were collected at day 2 and analyzed for cell viability (MTS assay). PFA-fixed slices used as negative control. Statistical analysis was calculated using two-way ANOVA analysis for statistical significance as mean ± SEM. * = p-value < 0.05.

Ciraku et. al., Extended Figure 1

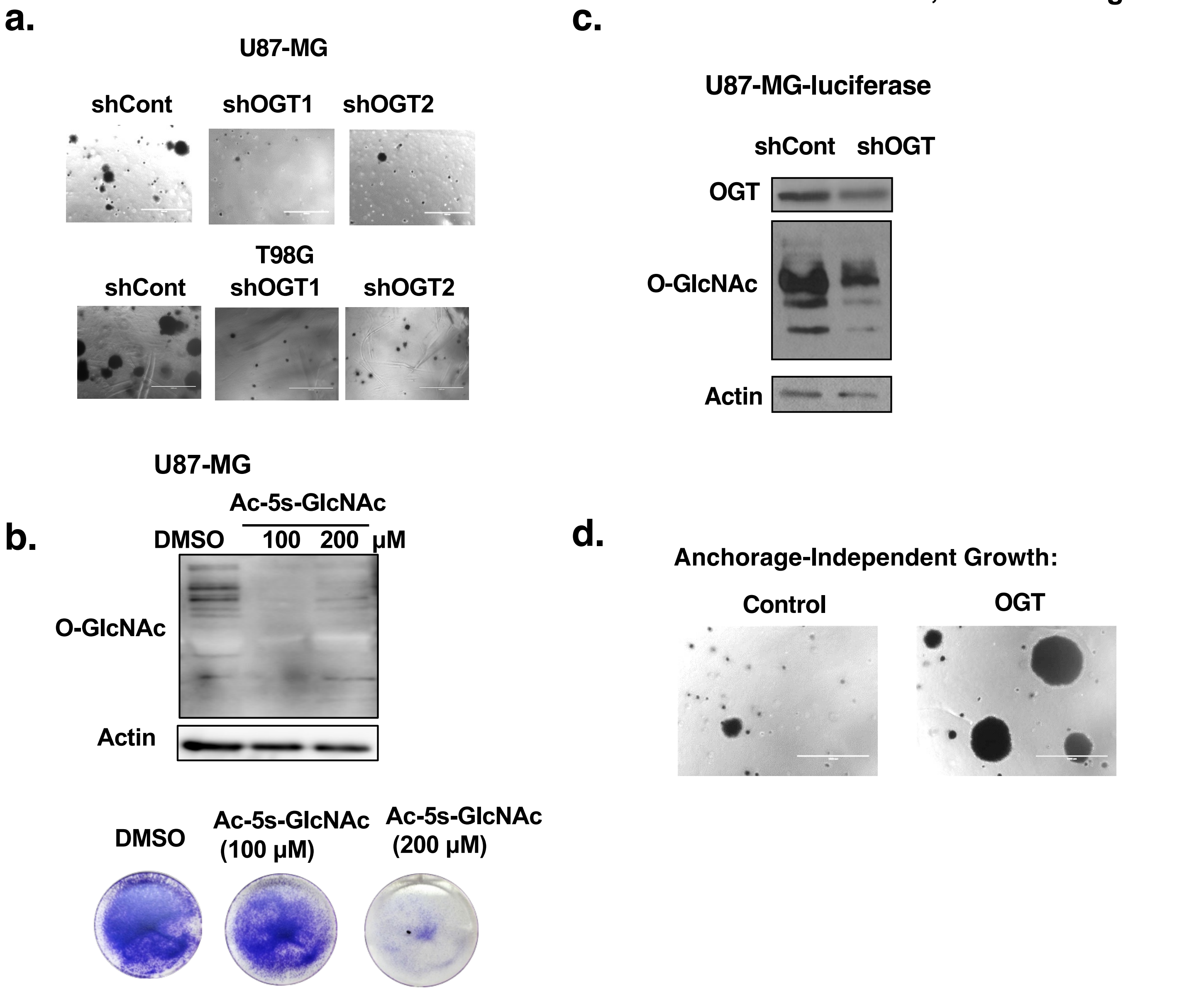

Ciraku, et. al., Extended Figure 2

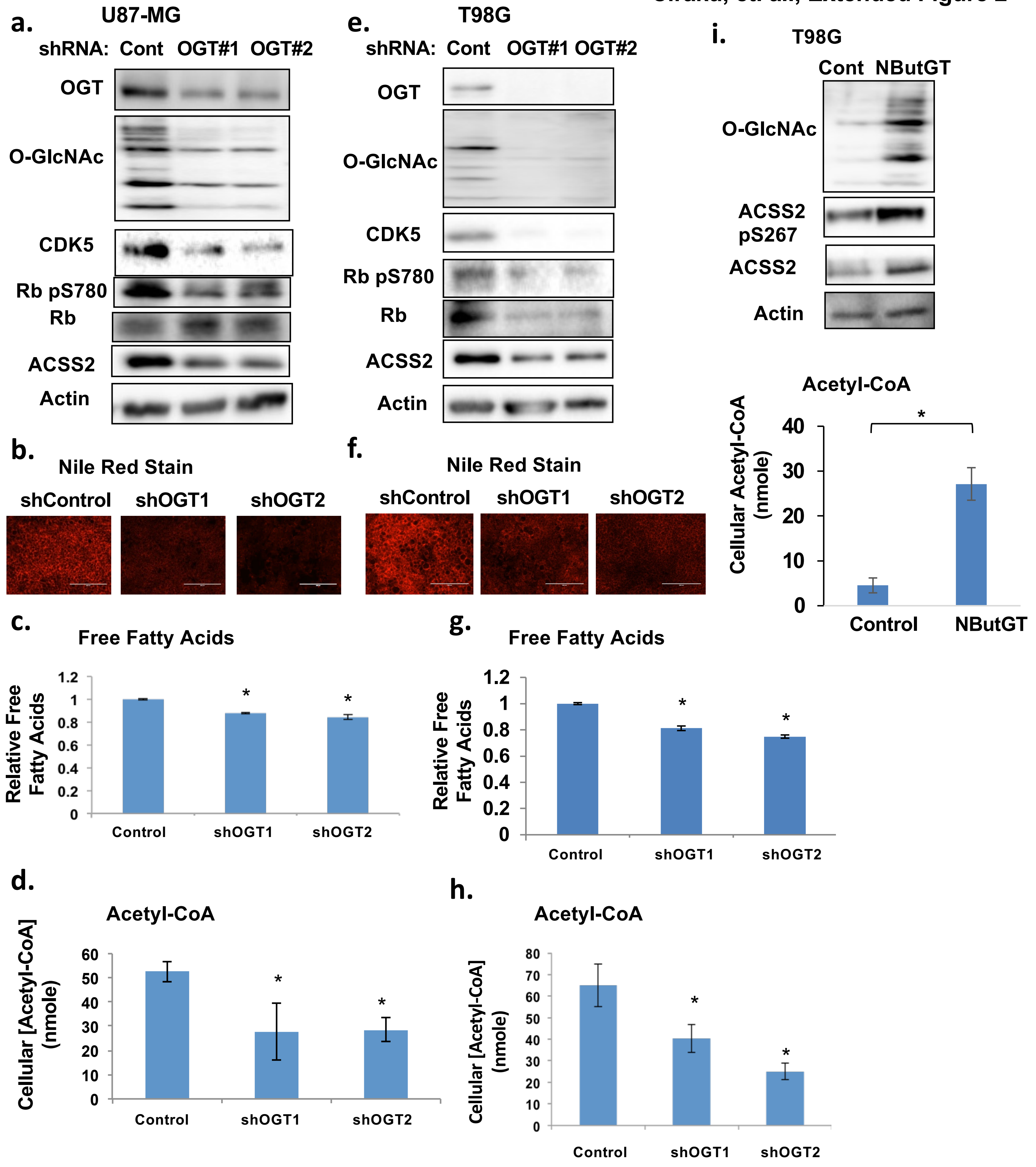

Ciraku, et. al., Extended Figure 3

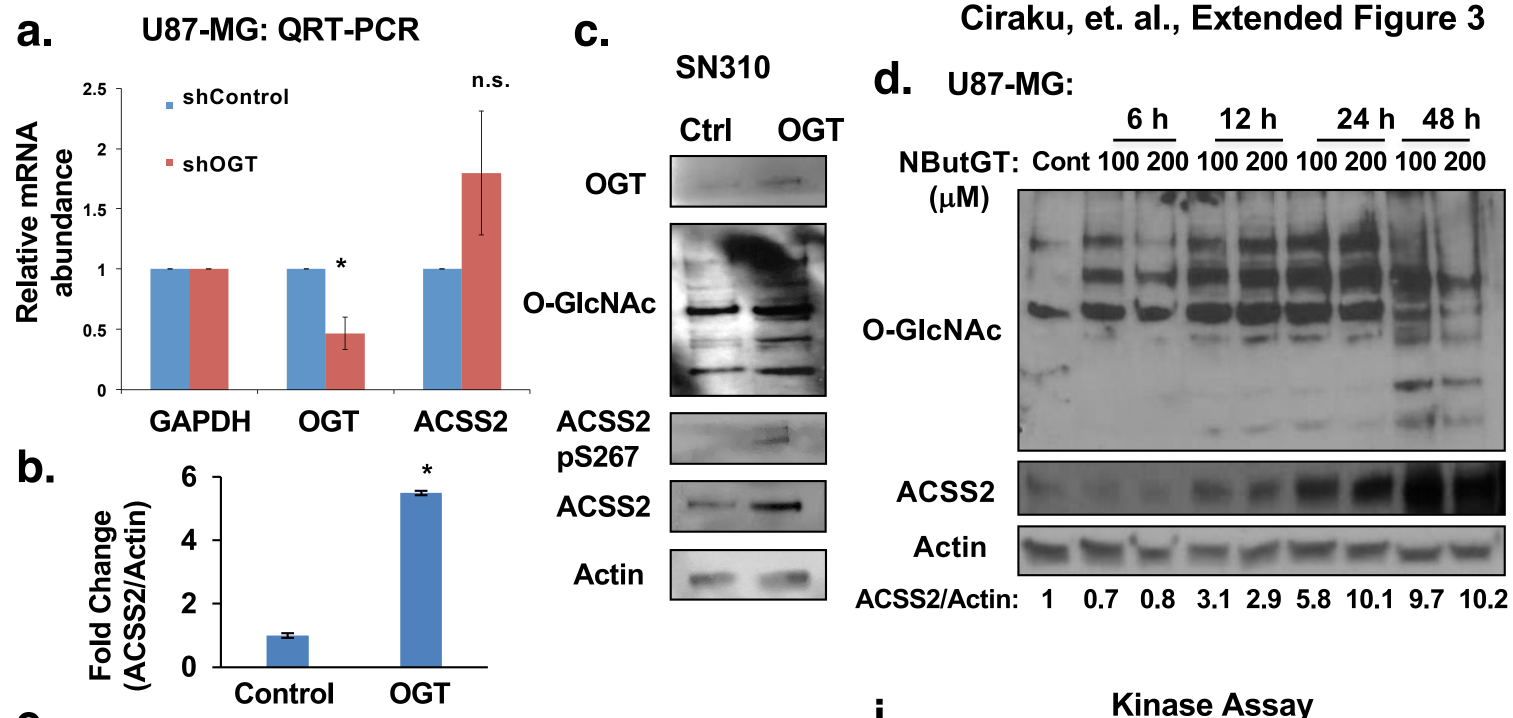


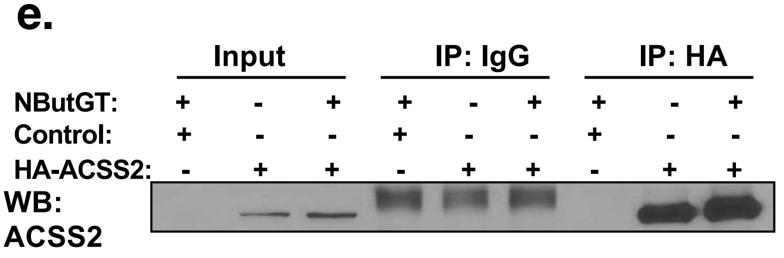


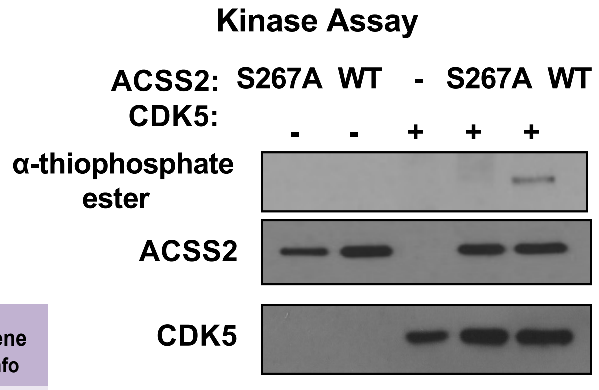


f.

| Score | Percentile | Motif | Motif Group | Site | Sequence | Surface Accessibility | Gene Info |
|---|---|---|---|---|---|---|---|
| 0.475 | 0.87% | CDK5 Kinase | Proline-dependent serine/threonine kinase group | S267 | STSQsPPIKR | 1.19 | CDK5 |

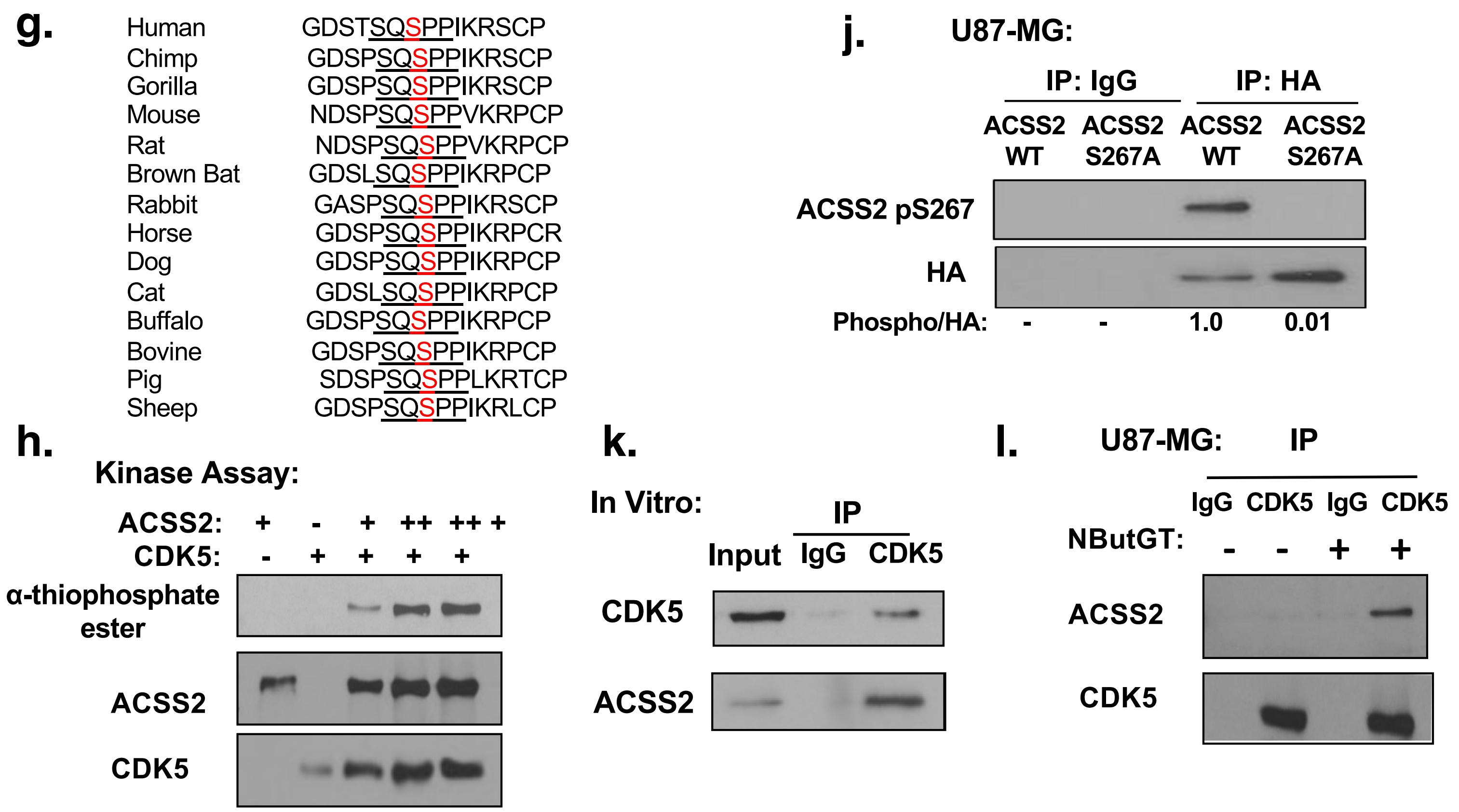

Ciraku et al. Extended Figure 4

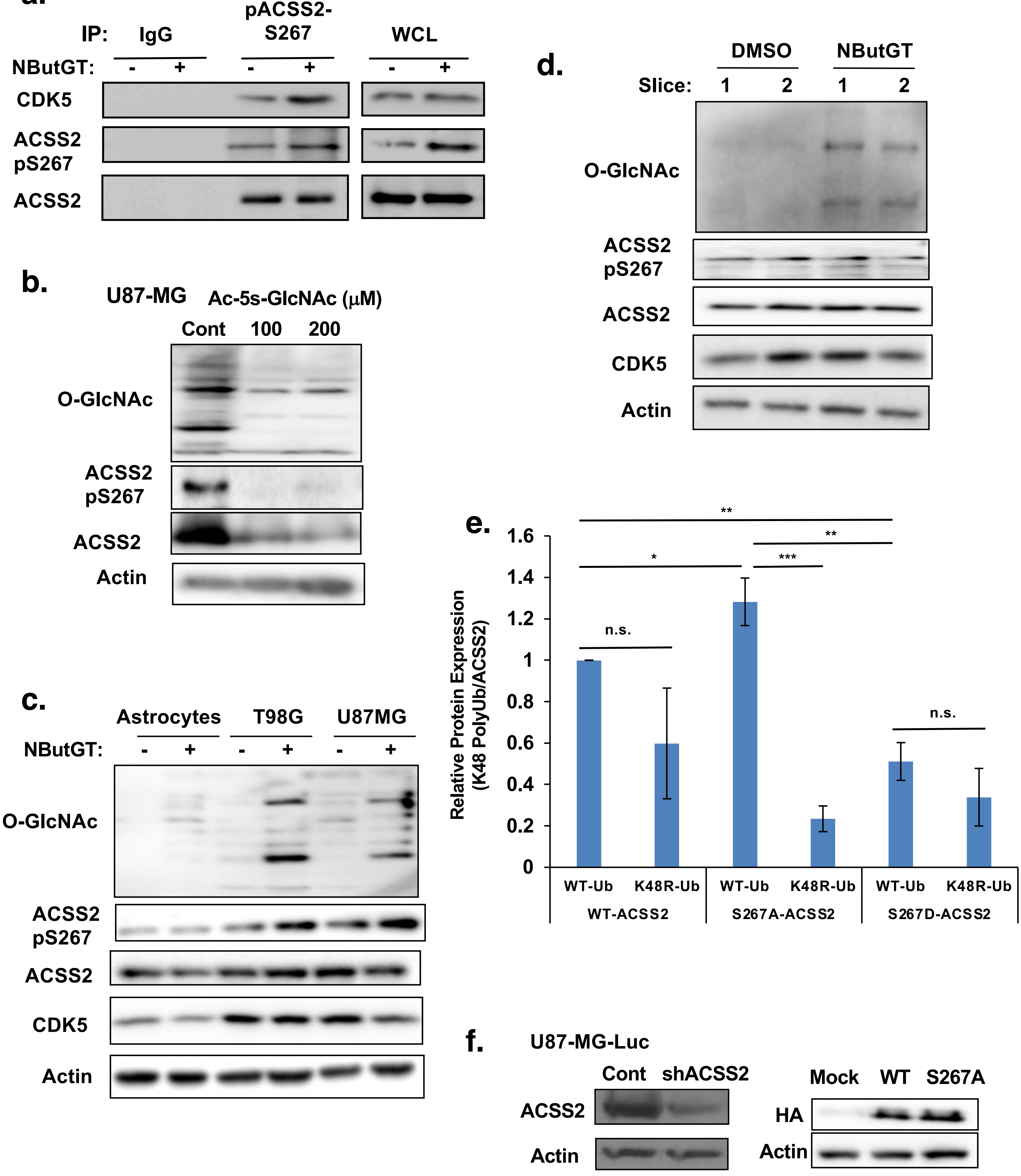

### Ciraku et al. Extended Figure 5

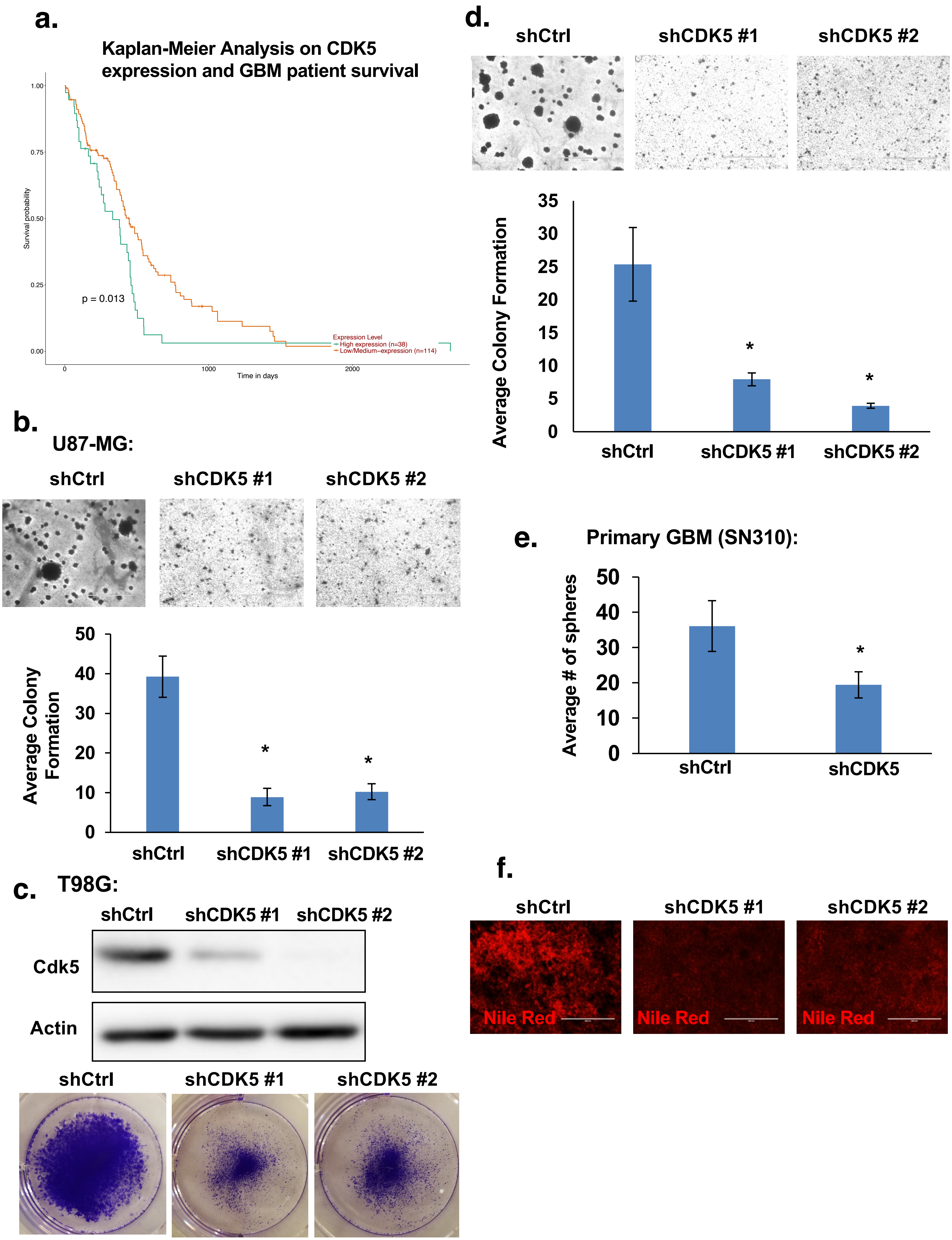

Ciraku et al. Extended Figure 6

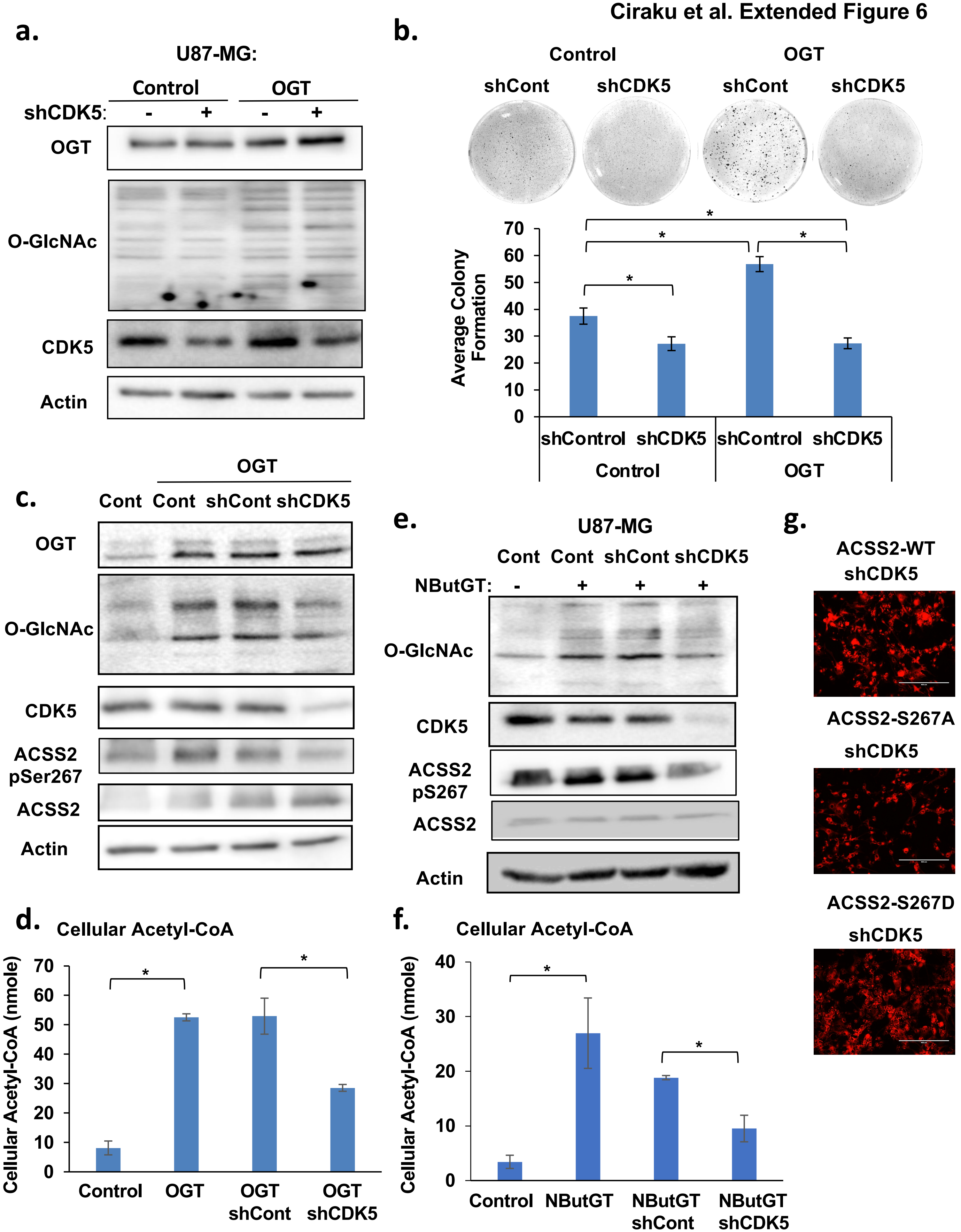

Ciraku et al. Extended Figure 7

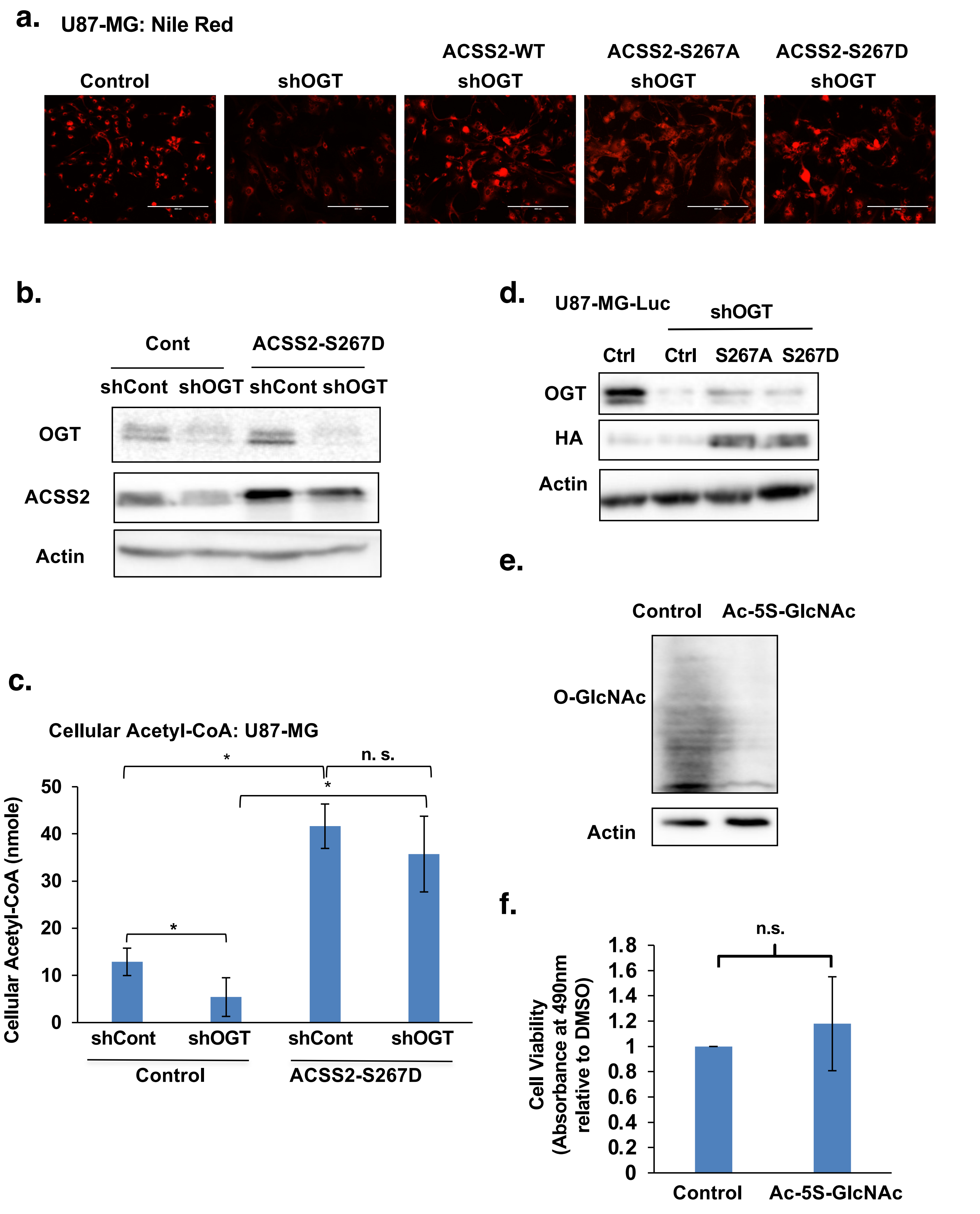

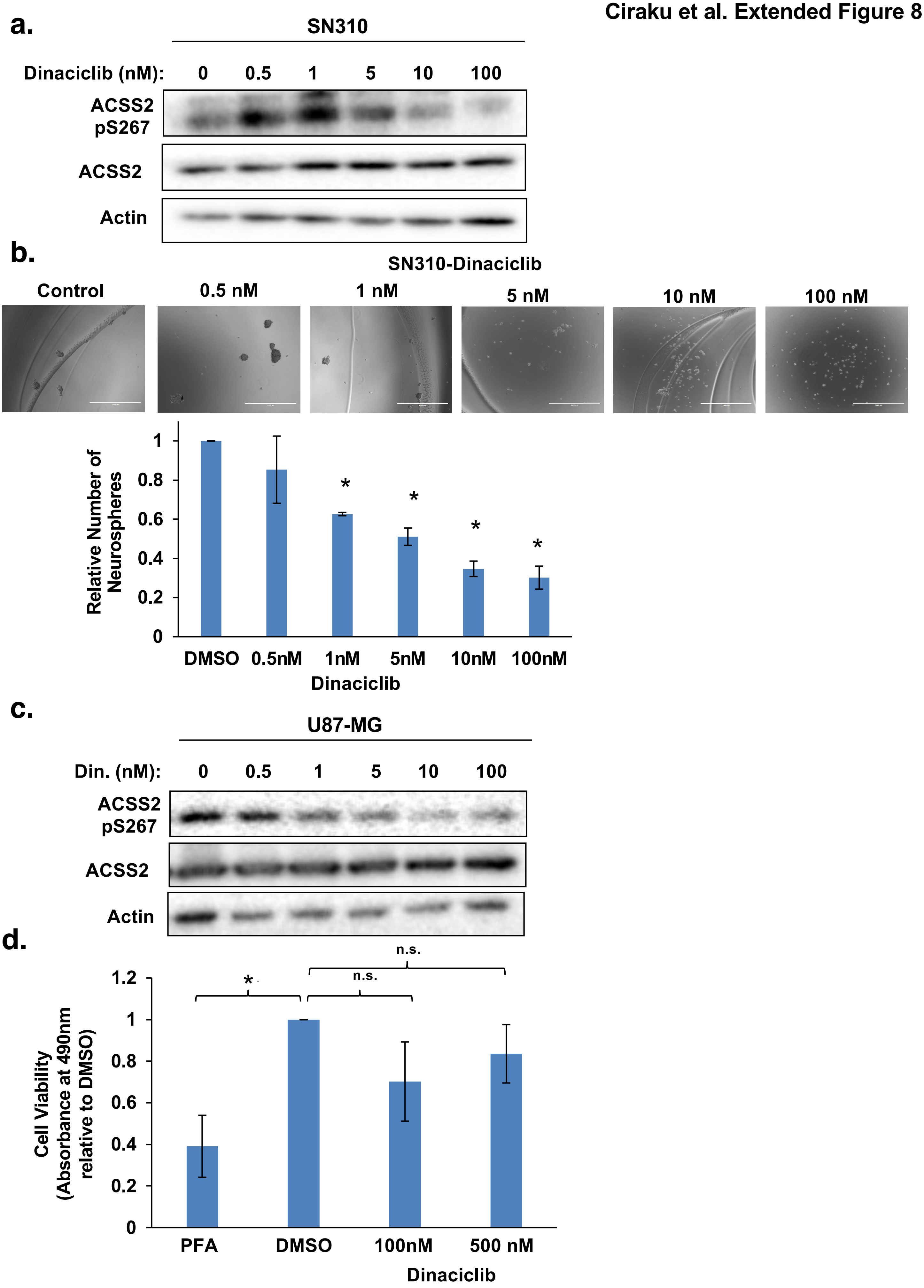
a.
SN310
Dinaciclib (nM): 0 0.5 1 5 10 100
ACSS2 pS267
ACSS2
Actin
b.
SN310-Dinaciclib
Control
0.5 nM
1 nM
5 nM
10 nM
100 nM
Relative Number of Neurospheres
0
0.2
0.4
0.6
0.8
1
DMSO 0.5nM 1nM 5nM 10nM 100nM
Dinaciclib
*
c.
U87-MG
Din. (nM): 0 0.5 1 5 10 100
ACSS2 pS267
ACSS2
Actin
d.
Cell Viability (Absorbance at 490nm relative to DMSO)
0
0.2
0.4
0.6
0.8
1
1.2
*
n.s.
n.s.
PFA DMSO 100nM 500 nM
Dinaciclib
